# Evolution of a putative, host-derived endosymbiont division ring and symbiosis-induced proteome rearrangements in the trypanosomatid *Angomonas deanei*

**DOI:** 10.1101/2021.08.23.457307

**Authors:** Jorge Morales, Georg Ehret, Gereon Poschmann, Tobias Reinicke, Lena Kröninger, Davide Zanini, Rebecca Wolters, Dhevi Kalyanaraman, Michael Krakovka, Kai Stühler, Eva C. M. Nowack

## Abstract

The transformation of endosymbiotic bacteria into genetically integrated organelles was central to eukaryote evolution. During organellogenesis, control over endosymbiont division, proteome composition, and physiology largely shifted from the endosymbiont to the host cell nucleus. However, to understand the order and timing of events underpinning organellogenesis novel model systems are required. The trypanosomatid *Angomonas deanei* contains a β-proteobacterial endosymbiont that divides synchronously with the host^1^, contributes essential metabolites to host cell metabolism^2-5^, and transferred one bacterial gene [encoding an ornithine cyclodeaminase (OCD)] to the nucleus^2^. However, the molecular mechanisms mediating the intricate host/symbiont interactions are largely unexplored. Here we identified seven nucleus-encoded proteins by protein mass spectrometry that are targeted to the endosymbiont. Expression of fluorescent fusion proteins revealed recruitment of these proteins to specific sites within the endosymbiont including its cytoplasm and a ring-shaped structure surrounding its division site. This structure remarkably resembles in shape and predicted functions mitochondrial and plastid division machineries. The endosymbiotic gene transfer-derived OCD localizes to glycosomes instead of being retargeted to the endosymbiont. Hence, scrutiny of protein re-localization patterns that are induced by endosymbiosis, yielded profound insights into how an endosymbiotic relationship can stabilize and deepen over time far beyond the level of metabolite exchange.

---

Besides the ancient endosymbiotic events that initiated the evolution of mitochondria and plastids more than one billion years ago, diverse bacterial lineages have evolved intimate endosymbiotic associations with eukaryotic hosts, often involving vertical endosymbiont transmission from one host generation to the next^6–8^. Similar to how eukaryotes control organelle abundance, a few protist hosts have additionally evolved the ability to strictly control the number of endosymbionts per host cell^1,9,10^. Over time, permanent host association results in gene losses and size reduction of endosymbiont genomes^11,12^. In these cases, the holobiont appears to rely on chimeric metabolic pathways involving enzymes encoded in both the endosymbiont and host genomes^2,13-16^. However, the molecular mechanisms enabling cross-compartment linkage of metabolic pathways, synchronization of host and endosymbiont cell cycles, and controlled segregation of endosymbionts to the daughter cells are largely unknown.

The most critical step in endosymbiont-to-organelle conversion supposedly is the evolution of a dedicated protein translocation system that enables the import of nucleus-encoded proteins into the endosymbiont^17^. The cases of the cercozoan amoeba *Paulinella* where hundreds of nucleus-encoded proteins are imported into the cyanobacterial endosymbiont^18^, and mealybug insects where peptidoglycan (PG) biosynthesis in the innermost of two nested bacterial endosymbionts depends on the import of a nucleus-encoded D-Ala-D-Ala ligase^19^, suggest that organellogenesis events are not restricted to mitochondria and plastids but can occur in more recently established endosymbiotic associations too. Also, in a few other systems with vertically transmitted bacterial endosymbionts, there are scattered reports on single host proteins that translocate into the endosymbiont cytoplasm^20,21^. Deciphering the rules that lead to the evolution of host control over a bacterial endosymbiont and endosymbiont-to-organelle transition would depend on the comprehensive proteomic characterization of further endosymbiotic associations and the development of efficient genetically tractable model systems for endosymbiosis.

The trypanosomatid *A. deanei* (subfamily Strigomonadinae) is an emerging model system to study endosymbiosis^22,23^. All members of the Strigomonadinae carry a β-proteobacterial endosymbiont (family Alcaligenaceae)^23,24^. *Candidatus* Kinetoplastibacterium crithidii, the endosymbiont of *A. deanei*, lies surrounded by a bacterial inner and outer membrane and a reduced PG layer free in the host cytosol^25^. Strict synchronization of the host and endosymbiont cell cycles results in a single endosymbiont per daughter cell after cell division^1^. The endosymbiont genome (0.8 Mbp) is highly streamlined and lost most genes for the core energy metabolism as well as the biosynthetic capacity for amino acids and cofactors such as proline, cysteine, and biotin; other biosynthetic pathways (e.g., for aromatic amino acids, riboflavin, and heme) were retained and apparently contribute to the host metabolism ^2,5,26,27^. To enable scrutiny of host/endosymbiont interactions, we previously developed genetic tools for *A. deanei* that allow for transgene expression and targeted gene knock-outs^28^. Furthermore, we identified one nucleus-encoded protein of unknown function, termed endosymbiont-targeted protein 1 (ETP1), that specifically localizes to the endosymbiont^28^, suggesting that protein targeting to the endosymbiont plays a role in host/endosymbiont interaction.

To determine the extent of protein import into *Ca*. K. crithidii, we analyzed proteins extracted from isolated endosymbionts (ES samples) (**Fig. 1a-b**) and whole cell lysates (WC samples) by liquid chromatography coupled to tandem mass spectrometry (LC-MS/MS). Two independent proteomic analyses totaling 9 biological replicates detected with high confidence overall 573 and 638 endosymbiont-encoded proteins (i.e., 78% and 87% of the 730 predicted endosymbiont-encoded proteins^26^) and 2,646 and 2,175 host-encoded proteins, respectively (**Supplementary Table S1**). Proteins identified exclusively or appearing enriched in ES samples comprised not only endosymbiont-encoded but also several host-encoded proteins (**Fig. 1c-d**).

**Fig. 1:**
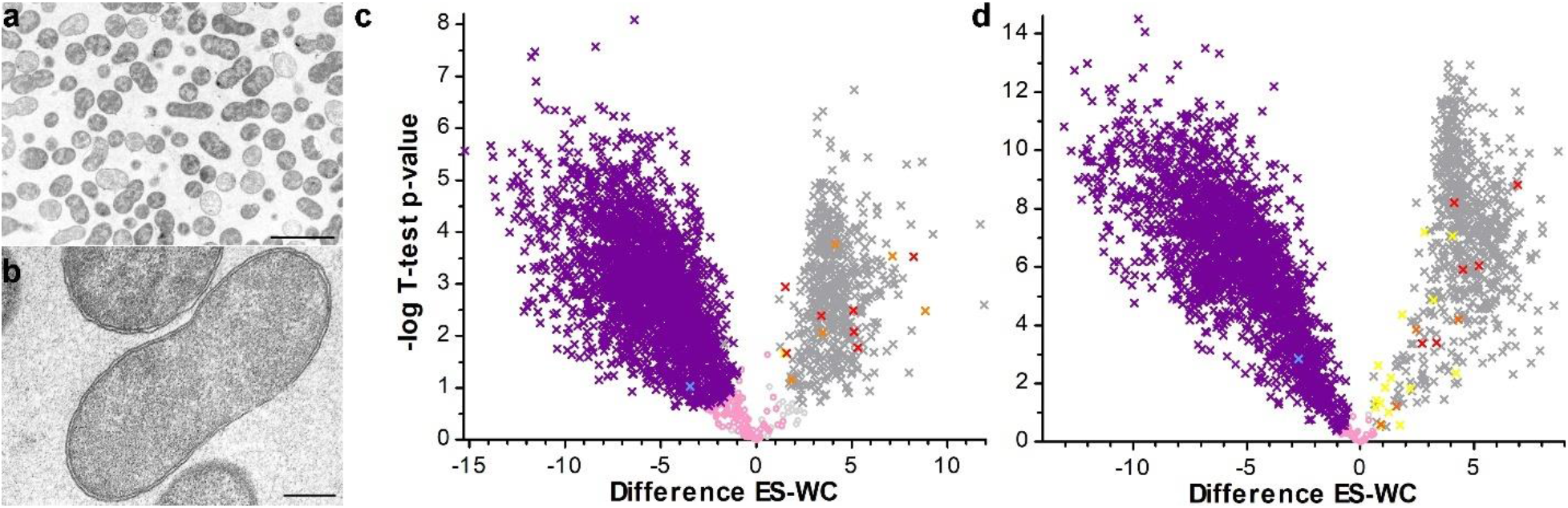
Comparative proteome analysis of whole cell lysates (WC) versus purified endosymbionts (ES) of *A. deanei*. **(a-b)** Transmission electron microscopy (TEM) of *Ca*. K. crithidii. **(a)** Overview of the collected endosymbiont fraction. Most of the structures observed in this fraction consist of eight-shaped and round structures that are surrounded by a double membrane consistent with the endosymbiont. Scale bar: 2.5 μm. **(b)** The endosymbiont outer and inner membrane remained intact in most of the cells during isolation. Scale bar: 250 nm. **(c, d)** Volcano plots of proteins identified by LC-MS/MS in Experiment 1 (**c**) and Experiment 2 (**d**). The difference of intensities of individual proteins between WC and ES samples 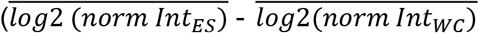; Difference ES-WC) is plotted against significance (-log10 p-value in Student’s T-test) for proteins detected in ES or WC samples. Color code: grey, endosymbiont-encoded proteins; colorful, nucleus-encoded proteins (red, enriched in endosymbiont in both experiments; orange, enriched in ES samples in one experiment, not identified in the other experiment; yellow, enriched in ES samples in one experiment, but depleted in the other experiment; blue, the endosymbiotic gene transfer (EGT)-derived OCD; purple, remaining nucleus-encoded proteins). Crosses in bright colors, significant enrichment or depletion; circles in pale colors represent nonsignificant values (rose, nucleus-encoded; light grey, endosymbiont-encoded).

Host-encoded proteins that either showed significant enrichment in the endosymbiont fraction in both experiments (red crosses in **Fig. 1c-d**) or showed significant enrichment in one experiment but were not detected at all or showed only nonsignificant enrichment in the endosymbiont fraction in the other experiment (orange crosses in **Fig. 1c-d**) were considered as putative endosymbiont-targeted proteins (ETPs). This group of 14 putative ETPs (for details see **Table S2**) also contained the previously identified ETP1^28^. Nucleus-encoded proteins that appeared as endosymbiont-enriched in one experiment but as host-enriched in the other experiment (yellow crosses in **Fig. 1c-d**) contained several predicted glycosomal or mitochondrial proteins. Thus, this group of proteins was regarded as putative contaminants and not further analyzed.

Next, we aimed to determine the subcellular localization of each of the 13 newly identified candidate ETPs in *A. deanei* using recombinant reporter protein fusions. We have previously demonstrated that ETP1 N- or C-terminally fused to the green fluorescent protein eGFP localized specifically to *Ca*. K. crithidii^*28*^. Therefore, a cell line background expressing ETP1 fused to the C-terminus of the red fluorescent protein mSCARLET (mS-ETP1) as an endosymbiont marker, was used for co-expression of the remaining 13 proteins of interest (POI) fused to the C- or N-terminus of eGFP (eGFP-POI and POI-eGFP, respectively). Western blot analyses of whole cell lysates and purified endosymbionts (up to the percoll step) obtained from the 13 cell lines co-expressing mS-ETP1 and each one of the eGFP-POI constructs showed that recombinant ETP1, ETP2, ETP3, ETP5, ETP7, and ETP8 are enriched in the endosymbiont fraction while recombinant ETP9 co-purifies to a certain extent with the endosymbiont (**Fig. 2a**). For the remaining candidate ETPs neither N-terminal nor C-terminal fusion constructs showed a signal in the endosymbiont fraction or (for one protein) no signal in the Western blot at all (**Supplementary Fig. 1**). Hence, these proteins were excluded from further analyses.

**Fig. 2:**
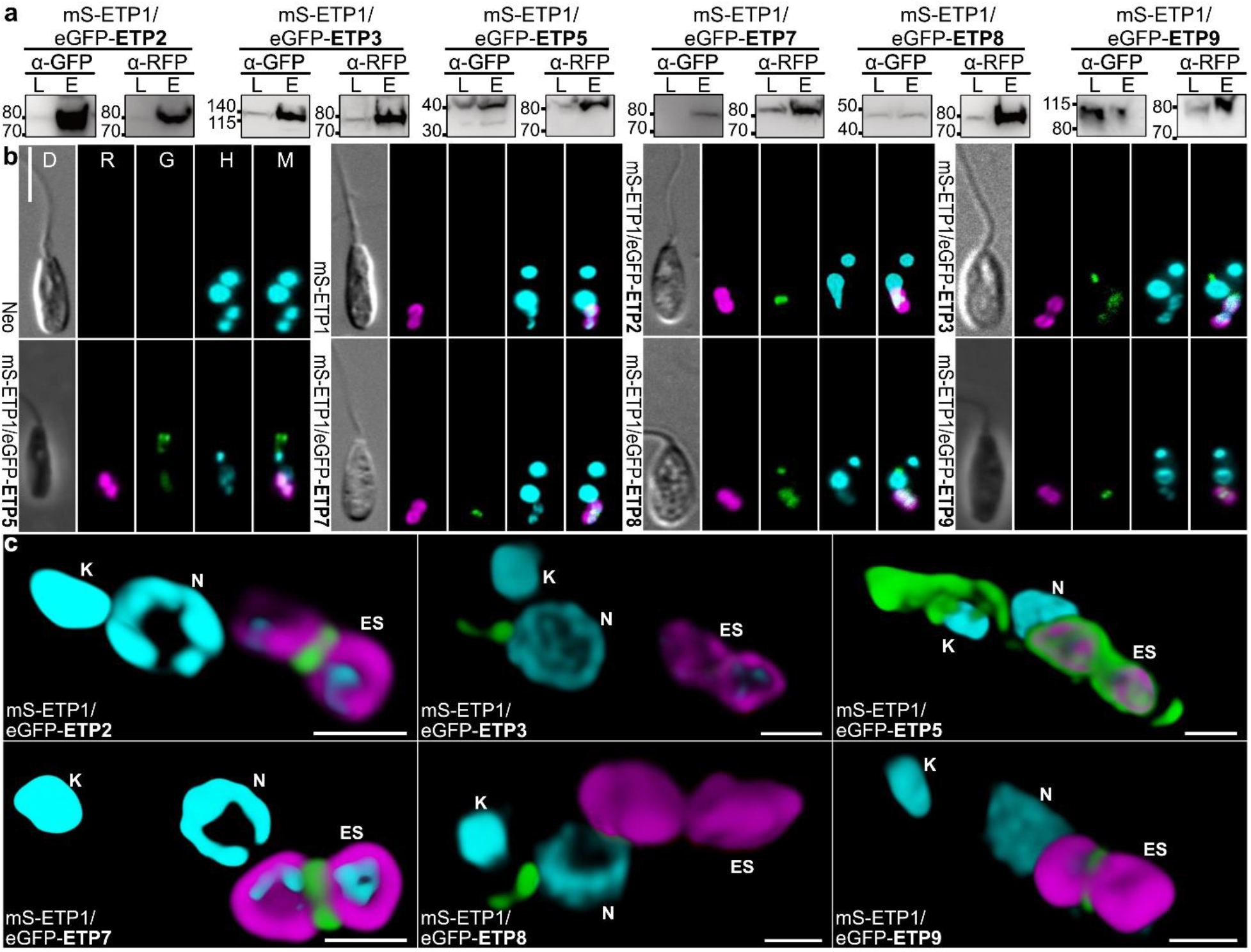
Newly identified ETPs show distinct subcellular localizations within *Ca*. K. crithidii and the host cell. (**a**) protein from whole cell lysate (L) or purified endosymbionts (E) were resolved by SDS-PAGE, transferred onto PVDF-membranes, and recombinant proteins visualized by Western blot analysis using anti-GFP (α-GFP) or anti-RFP (α-RFP) antibodies. (**b**) Epifluorescence microscopic analysis of cell lines expressing the neomycin phosphotransferase (Neo) alone, mS-ETP1, or mS-ETP1 in combination with eGFP-POI constructs. D, differential interference contrast; R, red channel; G, green channel; H, blue channel visualizing Hoechst 33342 staining; M, merge of the three fluorescence channels. Scale bar is 5 μm. **(c)** Three-dimensional reconstruction of the localization of the different recombinant ETPs within *A. deanei* from the superposition of 12-32 Z-stacks after deconvolution. Color code: magenta, mS-ETP1; green, eGFP-POI; cyan, Hoechst33342. Scale bar is 1 μm. ES, endosymbiont; K, kinetoplast; N, nucleus.

Localization of ETP2, ETP3, ETP5, ETP7, ETP8, and ETP9 at the endosymbiont was further confirmed by epifluorescence microscopy (**Fig. 2b**). Interestingly, the various ETPs localize to specific sites within the endosymbiont. Recombinant ETP2, ETP7, and ETP9 localize specifically at the constriction site of the eight-shaped endosymbiont; ETP5 localizes to the endosymbiont and the host cell flagellar pocket; ETP3 and ETP8 show a diffuse eGFP signal over the endosymbiont and additionally in a defined dot-like structure near the host cell nucleus (**Fig. 2b**). As previously observed for ETP1^28^, shifting the eGFP-tag to the C-terminal end of the ETPs did not affect the localization of ETP2, ETP5, ETP7, ETP8, and ETP9; only ETP3-eGFP did not yield a green fluorescence signal (**Supplementary Fig. 2**).

3D reconstruction of the fluorescence signal of each recombinant ETP obtained from focal planes of confocal fluorescence microscopy (**Fig. 2c** and **Supplementary Movies 1-6**) revealed that ETP1 is confined to the endosymbiont envelope; ETP2, ETP7, and ETP9 form a ring-shaped structure around the constriction site of the endosymbiont (i.e., the site where endosymbiont division occurs); ETP5 localizes at the host cell flagellar pocket from where thin fiber-like projections surround the periphery of the endosymbiont and seem to associate with the kinetoplast and nucleus of the host cell. ETP3 and ETP8 apparently localize inside the endosymbiont, indicating that these proteins translocate across the endosymbiont envelope membranes. Interestingly, ETP3 and ETP8 are additionally found in a barbell-shaped structure that sits on the anterior side of the nucleus. This structure is very similar in shape and positioning to the Golgi apparatus of *Trypanosoma brucei* and *Leishmania donovani*^29,30^. The Golgi is the main hub of vesicular trafficking in eukaryotic cells and, in different endosymbiotic associations, nucleus-encoded, endosymbiont-targeted proteins traffic through the Golgi^21,31-33^. However, an important difference in these associations is that the outermost membrane surrounding the endosymbiont is host-derived. Nevertheless, vesicles that appear to fuse with the outer endosymbiont membrane have been observed before in electron micrographs of Strigomonadinae^34^, raising the possibility that Golgi-derived vesicles can target *Ca*. K. crithidii in *A. deanei*. However, none of the ETPs contain a predicted targeting signal for the secretory pathway (nor the mitochondrion), all are soluble, and comparison of the ETP protein sequences among each other did not reveal any obvious common characteristics such as similar sequence extensions that could serve as targeting signals or common motifs (as analyzed by MEME 5.0.5 ^35^; cut-off: e-value <0.05).

To explore the cellular functions of the ETPs, we aimed to generate null mutants of ETP1, ETP2, and ETP7. However, whereas heterozygous knock-out mutants could be obtained, deletion of both alleles of the corresponding genes did not yield viable clones in all cases after several attempts. The inability to generate homozygous knock-out mutants is suggesting an essential function of these proteins. Since no inducible gene expression systems are available yet for *A. deanei*, a functional characterization of these genes is yet to come. Nevertheless, for several ETPs, observed subcellular localizations and functional annotations (**Table 1**) imply their involvement in distinct cellular processes.

**Table 1:** Endosymbiont-targeted proteins in *A. deanei*.

| ETP | Annotation | Best Blastp Hit<sup>a</sup> | e-value | %id<sup>b</sup> | BBH<sup>c</sup> |
| --- | --- | --- | --- | --- | --- |
| ETP1 | Hypothetical protein, cons. | - | - | - | - |
| ETP2 | Hypothetical protein, cons. | - | - | - | - |
| ETP3 | Hypothetical protein, cons. | hypothetical protein<br>JIQ42_00906 [*Leishmania* sp.<br>Namibia] | 2e-19 | 26 | No |
| ETP5 | Kinetoplastid membrane protein<br>11, put. | kinetoplastid membrane protein<br>KMP-11 [*Trypanosoma cruzi*<br>strain CL Brener] | 7e-52 | 94 | Yes |
| ETP7 | Phage tail lysozyme, put. | - | - | - | - |
| ETP8 | Hypothetical protein, cons. | unnamed protein product<br>[*Phytomonas* sp. isolate EM1] | 3e-12 | 28 | Yes |
| ETP9 | Dynamin family/Dynamin central<br>region/Dynamin GTPase effector<br>domain containing protein,<br>putative | unnamed protein product<br>[*Trypanosoma congolense*<br>IL3000] | 1e-114 | 34 | No |

<sup>a</sup> Protein with the lowest e-value (outside *A. deanei*) returned by Blastp against the NCBI nr database as of July 14, 2021 (e-value cut-off of 1e-6).
<sup>b</sup> Percentage of amino acid identity between best Blastp hit and the corresponding ETP.
<sup>c</sup> Best bidirectional blast hits obtained between the NCBI nr database and an in-house database containing the previously generated *A. deanei* transcriptome.

The ring-shaped arrangement of ETP2, ETP7, and ETP9 around the endosymbiont division site, is suggestive of a function in endosymbiont division. For all three recombinant proteins, this ring structure is seen in only around half of the cells in a mid-log phase culture (**Supplementary Fig. 3a**). *Ca*. K. crithidii encodes FtsZ, a GTPase that typically self-assembles into a ring structure at the inner side of the cytoplasmic membrane at bacterial division sites initiating cytokinesis, and the Min system which is typically involved in positioning the FtsZ ring^36^. However, antibodies specific against FtsZ distribute evenly throughout the cell instead of localizing to a division ring^37^, suggesting that the bacterial division machinery might not be fully functional in *Ca*. K. crithidii. Furthermore, exposure of *A. deanei* to the eukaryotic translation inhibitor cycloheximide not only results in cessation of host cell growth but also blocks endosymbiont division^38^, suggesting the involvement of host-derived factors in endosymbiont division. Intriguingly, ETP9 is annotated as ‘dynamin family protein’. Members of the dynamin family are self-assembling, polymer-forming GTPases that are involved in diverse cellular membrane remodeling events. In the Opisthokonta, three dynamin-related proteins (DRPs) are involved in the dynamic fission and fusion of mitochondria^39^. Assembly of the soluble cytosolic DRP, DNM1/DRP1 (in yeast/human), into helical oligomers on the mitochondrial membrane and constriction upon GTP hydrolysis leads to mitochondrial fission^40,41^. Trypanosomatids outside the Strigomonadinae encode only a single DRP (or, in *T. brucei*, two nearly identical, functionally likely equal, tandemly duplicated DRPs)^42,43^. This ‘dynamin-like protein of *T. brucei*’ (TbDLP) shows high homology to DNM1 in yeast and was shown to regulate mitochondrion division as well as endocytosis^42,43^. Interestingly, in *A. deanei* there are two divergent DLPs, AdDLP (CAD2218610.1) and ETP9 (CAD2212698.1). AdDLP shows 68% identity to TbDLP; ETP9 contains the N-terminal GTPase and C-terminal GTPase effector domain typical for DRPs but shares only 34% identity with TbDLP (**Supplementary Fig. 3b**). The *etp9* gene might have evolved by duplication and divergence of *Addlp* but localizes on a different chromosome. Similarly, also in the Archaeplastida which acquired a cyanobacterial endosymbiont that evolved into the plastid, a plant-specific dynamin evolved-likely by duplication and divergence of a DRP involved in cytokinesis^44^. Upon plastid division, this plant-specific dynamin is recruited to the plastid division site and forms a constriction ring on the cytosolic surface of the outer membrane that seems to aid with constriction and mediates the final fission of the plastid^45^.

ETP7 is annotated as ‘phage tale lysozyme’. Phyre2^46^ predicts with 92.6% confidence structural homology of the C-terminal part of ETP7 (aa 358-518) with a cell wall degrading enzyme in the bacteriophage φ29 tail^47^. Although sequence identity is only 24% between both proteins, catalytic site and PG-binding site seem to be conserved in ETP7. Intriguingly, PG hydrolysis by a nucleus-encoded enzyme that localizes at the plastid division site is essential also for plastid division in the Glaucophyte algae that possess a PG layer between the two envelope membranes and at least in some basally branching Viridiplantae^48^. This finding suggests that during the early stages of plastid evolution, the ancestral algae regulated plastid division by PG splitting by a nucleus-encoded enzyme.

The function of ETP2, that shows neither amino acid sequence nor structural homology to known proteins and is predicted to be mainly unstructured, remains unclear. However, its apparent arrangement in the putative endosymbiont division ring suggests its involvement in symbiont division. Also the plastid and mitochondrial division machinery contain besides dynamin-related proteins and (sometimes) PG hydrolases other proteins. These have various functions, such as recruitment of soluble factors to the organelle membranes or cross-membrane linkage of the bacterium-derived and host-derived components of the organelle division machinery^49–51^.

ETP5 shows high sequence identity (94%) to the ‘kinetoplastid membrane protein 11’ (KMP-11) of *T. cruzi* which is highly conserved across trypanosomatids^52^. As in other trypanosomatids, ETP5 is encoded by a multicopy gene and occurs in four tandemly arranged identical gene copies. In *T. brucei, T. cruzi*, and *Leishmania infantum*, KMP-11 localizes to the basal body, flagellar pocket, and flagellum^53,54^, and associates with microtubules^55^. Although its exact cellular function is unknown, its depletion blocks cytokinesis in *T. brucei*^56^. The localization of ETP5 suggests that in *A. deanei* there is a host-derived structure that connects the three DNA-containing compartments (nucleus, mitochondrion, and endosymbiont) with the basal body. ETP5 likely interacts with these three compartments by direct interaction with the lipids present in their outer membranes^57^. Importantly, in trypanosomes, division of the basal body marks the transition from the G phase to the S phase in the cell cycle^58^. The basal body is physically linked to the kinetoplast through the tripartite attachment complex facilitating positioning and segregation of the replicated mitochondrial genome^59^. Thus, the observed localization of recombinant ETP5 in *A. deanei*, in combination with the cytokinesis defects following the silencing of the ETP5 ortholog KMP-11 in *T. brucei*^56^, suggests that ETP5 plays a role in orchestrating segregation of organelles and cellular structures during cytokinesis.

ETP1, ETP3, and ETP8 are annotated as hypothetical proteins. Blastp searches against the NCBI non-redundant (nr) database returned either no similar proteins from other organisms (for ETP1) or exclusively proteins of unknown function for ETP3 and ETP8 (**Table 1**). 3D structure prediction using Phyre2 revealed either no significant similarities to any known protein structures (for ETP1 and ETP8) or, with confidence levels >96%, similarity to several long stretched α-helical proteins with diverse functions such as muscle contraction, PG hydrolysis, or chromosome maintenance. Hence, predicting the cellular function of these proteins is impossible based on the data at hand.

Finally, the EGT-derived OCD (blue cross in **Fig. 1c,d**) was not among the candidate ETPs. Examination of the OCD amino acid sequence revealed that following EGT, the protein acquired a C-terminal peroxisomal targeting sequence type 1 (PTS1) in all members of the Strigomonadinae (**Fig. 3a**), suggesting that the protein localizes to the glycosome, a specialized peroxisome in trypanosomatids characterized by the presence of the first six or seven steps of glycolysis. Interestingly, glycosomes closely associate with the endosymbiont in the Strigomonadinae^22,60^ (**Fig. 3b**). Expression of a recombinant protein in which the OCD was fused to the C-terminus of eGFP (eGFP-OCD) in the background of a cell line expressing the glycosome-targeted red fluorescent protein mCHERRY-SKL, showed clear co-localization of eGFP-OCD with mCHERRY-SKL in epifluorescence microscopy confirming a glycosomal localization of the recombinant OCD (**Fig. 3d**). The OCD catalyzes the conversion of ornithine to proline. Ornithine can be formed in the glycosome by the activity of the arginase, which also contains a PTS1 and is localized in the glycosomes in *Leishmania* ssp.. Thus, re-localization of the OCD to the glycosome likely results in proline production within the glycosome. Since *Ca*. K. crithidii lost the ability to generate proline, which is required for protein biosynthesis in the endosymbiont, the observed close proximity of the endosymbiont to proline-generating glycosomes might support the metabolic integration of the endosymbiont in *A. deanei* (**Fig. 3c**).

**Fig. 3:**
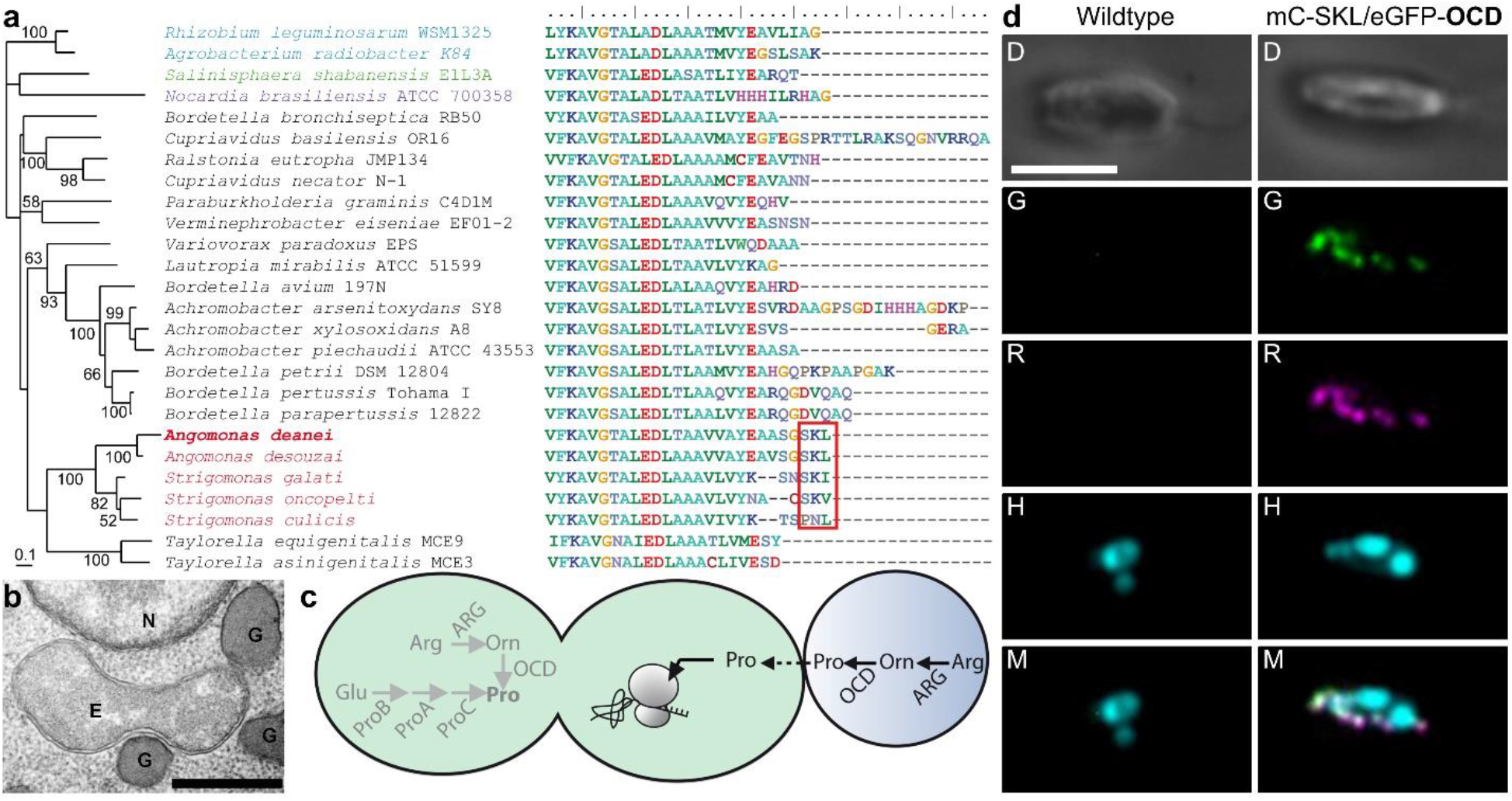
The EGT-derived OCD in the Strigomonadinae acquired a PTS1 signal and localizes to the glycosome. **(a)** Left, maximum likelihood phylogeny of the OCD of bacteria and the Strigomonadinae (taxon sampling according to ref.^2^). Species names are colored according to taxonomic affiliation. Red, Stigomonadinae; black, β-proteobacteria; green, γ-proteobacteria; blue, α-proteobacteria; violet, actinobacteria. Values at branches represent bootstrap support >50%. Right, alignment of the C-termini of the corresponding proteins. Red box, PTS1. **(b)** TEM of *A. deanei* shows *Ca*. K. crithidii surrounded by several glycosomes. E, endosymbiont; G, glycosome; N, nucleus; scale bar is 500 nm. **(c)** Scheme of proline metabolism in *A. deanei*. Endosymbiont, green; glycosome, blue. Arrows in grey represent enzymes missing from the endosymbiont genome; arrows in black, enzymes encoded in the nuclear genome. Dashed arrow represents metabolite transport. ARG, arginase (EC:3.5.3.1); OCD, ornithine cyclodeaminase (EC:4.3.1.12); ProB, glutamate 5-kinase (EC:2.7.2.11); ProA, glutamate-5-semialdehyde dehydrogenase (EC:1.2.1.41); ProC, pyrroline-5-carboxylate reductase (EC:1.5.1.2). **(d)** Epifluorescence microscopic analysis of *A. deanei* cell line co-expressing eGFP-OCD and mCherry-SKL. Fluorescence channels are as in Fig. 2b,c. Scale bar: 5 μm.

In sum, we identified seven ETPs in *A. deanei* representing a combination of typical trypanosomatid proteins and proteins that evolved newly in *A. deanei* (or diverged beyond recognition from their original source). Their discrete subcellular localizations within the endosymbiont, as well as their functional annotations, suggests their involvement in distinct biological processes. We postulate that, convergent to the evolution of the plastid division machinery, a dynamin and PG hydrolase-based host-derived division ring system evolved, that provides *A. deanei* with control over the division of its endosymbiont. Despite the apparent capacity of specific nucleus-encoded proteins (ETP3 and ETP8) to translocate across the endosymbiont membranes, cross-compartment linkage of metabolic pathways seems to rely rather on metabolite shuttling than protein import. Metabolic integration of the endosymbiont might be facilitated by the tight association of glycosomes that produce metabolites required by the endosymbiont. In conclusion, our work demonstrates that in addition to studying gene presence/absence patterns by genomics, analysis of symbiosis-induced protein re-localization, is key to understand the molecular mechanisms guiding endosymbiotic interactions. The results obtained strongly support the emerging pattern that protein import evolves early during endosymbiosis providing the host with control over the endosymbiont^7^ and not as a consequence of EGT to enable re-import of the gene products of the transferred genes.

## Methods

### Culture conditions and generation of transgenic cell lines

*Angomonas deanei* (ATCC PRA-265) was grown as described before^28^. For transfection, between 1 x 10^6^ and 1 x 10^7^ cells were resuspended in 18 μl of P3 primary cells solution (Lonza), 2 μl of the restricted cassette (2-4 μg total) were added, and cells were pulsed with the program FP-158 using the Nucleofector 4D (Lonza). After transfection, cells were transferred to 5 ml of fresh brain-heart infusion medium (BHI, Sigma Aldrich) supplemented with 10% v/v horse serum (Sigma Aldrich) and 10 μg/ml of hemin, incubated at 28 °C for 6 h, and then diluted 10-fold in the same media containing the selection drug(s) (i.e., G418 at 500 μg/ml, hygromycin B at 500 μg/ml, and/or phleomycin at 100 μg/ml final concentration). Aliquots of 200 μl were distributed onto 96-well plates and incubated at 28 °C until clonal cell lines were recovered, typically between 5-7 days. Correct insertion of the cassette was verified by PCR.

### Isolation of the endosymbiont of *A. deanei* and proteomic analysis

Endosymbionts were isolated from *A. deanei* cells lysed by sonication on consecutive sucrose, percoll, and iodixanol gradient as described previously^28^. Finally, endosymbionts were resuspended in 200 μl of buffer B (25 mM Tris-HCl, pH 7.5, 20 mM KCl, 2 mM EDTA, and 250 mM sucrose). The resulting endosymbiont fractions mainly consisted of intact endosymbionts as judged by the presence of a double membrane surrounding the endosymbionts viewed by TEM (**Fig. 1a-b**) and were thus considered suitable for proteomic analyses. Proteins from isolated endosymbionts were either precipitated by addition of trichloroacetic acid (TCA) to a final concentration of 10% v/v, washed 2x in cold acetone, and resuspended in 200 μl 0.1 N NaOH (Experiment 1) or the isolated endosymbionts were directly frozen in liquid nitrogen and stored at −80 °C until use (Experiment 2).

### Transmission electron microscopy

Isolated endosymbionts obtained from the iodixanol gradient were fixed overnight in 2.5% glutaraldehyde in 0.1 M cacodylate buffer, pH 7.4 at 4 °C. Fixed endosymbionts were pelleted at 7,600 x *g* for 5 min and resuspended in buffer containing 25 mM Tris-HCl, pH 7.5, 0.4 M sucrose, 20 mM potassium chloride, 2 mM ethylenediaminetetraacetic acid (EDTA), and 20% bovine serum albumin (BSA), incubated for 10 min on ice, and pelleted again. The resulting pellet was covered with 2.5% glutaraldehyde in 0.1 M cacodylate buffer taking care not to disrupt its integrity and fixed overnight at 4 °C. *A. deanei* cells grown to late exponential phase were washed twice with phosphate-buffered saline (PBS), fixed as described for isolated endosymbionts, and pelleted at 2,000 x *g* for 5 min. Pellets of fixed *A. deanei* cells and endosymbionts were washed once with 0.1 M cacodylate buffer, post-fixed with 2% osmium tetroxide plus 0.8% tetrasodium hexacyanoferrate, washed once more in 0.1 M cacodylate buffer, and embedded in 3.5% agar. Agar blocks containing the pellets were dehydrated by a graded series of ethanol from 60% to 100% and infiltrated with Epon-Aldarite using propylenoxide as an intermediate solvent. The resin was polymerized for 24 h at 40 °C and then 24 h at 60 °C. Thin sections of 70 nm were stained with lead citrate and uranyl acetate and examined with a transmission electron microscope (Zeiss 902) at 80 kV.

### Proteome analysis and identification of ETPs

Sample separation and LC-MS/MS analyses of WC and ES samples were essentially done as described before^28^. In total, the results from 9 biological replicates were analyzed. The first three biological replicates (Experiment 1) were run together in a preliminary analysis to check the quality of the endosymbiont preparation and differed from the later six biological replicates (Experiment 2) only by a TCA precipitation step of both the WC and ES samples. Peptides of tryptic digested samples were separated over 2h on C18 material using an Ultimate3000 rapid separation system (Thermo Fisher Scientific) as described^61^ and subsequently analyzed with an online coupled mass spectrometer in data dependent mode. Samples of Experiment 1 were analyzed using a QExactive plus mass spectrometer (Thermo Fisher Scientific) as described^61^ and samples of Experiment 2 analyzed on an Orbitrap Elite (Thermo Fisher Scientific) as described^62^.

Database searches were carried out with MaxQuant version 1.6.12.0 (MPI for Biochemistry, Planegg, Germany) using label-free quantification separately for the two analyzed groups (WC and ES) and standard parameters if not indicated otherwise. The ‘match between runs’ function was enabled, as well as LFQ and iBAQ^63^ quantification; LFQ quantification was carried out separately for WC and ES samples. Protein sequences as basis for searches were retrieved from UniProtKB (750 sequence entries from *Ca*. K. crithidii, downloaded on 9^th^ April 2019) and NCBI (10365 entries from GCA_903995115.1, *A. deanei*, downloaded on 1^st^ December 2020). The mass spectrometry proteomics data have been deposited to the ProteomeXchange Consortium via the PRIDE^64^ partner repository with the dataset identifier PXD017908.

Only proteins identified with at least 3 (Experiment 1) or 5 (Experiment 2) valid iBAQ intensities^65^, 2 different peptides, and at least 5% sequence coverage were considered as identified with high confidence and used in downstream analyses. Next, iBAQ intensities were normalized by dividing through the median iBAQ intensity of all proteins from the respective sample. For determination of endosymbiont-encoded proteins enriched in the host cell, missing values were imputed with values drawn from a downshifted normal distribution (downshift of 1.8 standard deviations, width 0.3. standard deviations) and two-sided Student’s t-test were calculated between normalized iBAQ values from WC and ES samples using the significance analysis of microarrays method^66^ to control for multiple testing (S0=0.6, false discovery rate 5%).

To verify correct prediction of translation start sites of each POI in the newly released, annotated *A. deanei* genome assembly (GCA_903995115.1)^67^ that was used as a database for mass spectrometric protein identification, the gene model of each POI was compared to the corresponding transcript in a previously generated *A. deanei* transcriptome dataset^28^. The longest possible N-terminal extension of the open reading frame (ORF) in 5’ full-length transcripts (as indicated by the presence of a 5’ splice leader sequence (SL)) was regarded as full-length ORF and used for further analyses (**Table S2, Supplementary Fig. 4**). Except for the hypothetic protein CAD2216283.1, all candidate ETPs were represented by transcripts with a full-length 5’ end. For this candidate, 5’ RACE allowed for extension of the transcript sequence up to the SL.

### RNA extraction, cDNA synthesis, and rapid amplification of cDNA ends (RACE)

Cells from 0.5 ml *A. deanei* cultures grown to late-logarithmic phase were collected by centrifugation, the pellet was frozen in liquid nitrogen and immediately resuspended in 1 ml of TRI Reagent (Sigma Aldrich). RNA was extracted according to the manufacturer’s instructions. RNA concentration was estimated by measuring the absorbance at 260 nm in a NanoDrop spectrophotometer (Thermo). 5 U of DNAase (Thermo) were added to 5 μg RNA, incubated for 10 min at room temperature to degrade residual DNA contamination, and DNase-treated RNA was purified using the RNAse MinElute Kit (Qiagen) according to the manufacturer’s instructions. 3 μg of DNase-treated RNA were used per RACE reaction using the 5’ RACE System for Rapid Amplification of cDNA Ends, version 2.0 (Thermo) with internal primers described in **Supplementary Table 3**. The obtained PCR fragments were cloned into the pJET 1.2 cloning vector (Invitrogen) and sequenced using the pJet Fw/Rv primer set provided by the manufacturer.

### Construction of plasmids

To efficiently generate eGFP-POI and POI-eGFP expression vectors for *A. deanei*, the pAdea043 and pAdea235 tagging vectors, respectively, were constructed (**Supplementary Fig. 5**). These plasmids target the insertion of the respective expression cassettes into the δ-amastin locus of *A. deanei*^28^. To this end, the *lacZ*α expression cassette encoding the alpha-fragment of the β-galactosidase under control of the lac promotor and operator was amplified from the circularized plasmid pGEM-T (Promega) using the forward primer 596 that includes a 5’ *Xho*I restriction site and the reverse primer 597 which includes a 3’ *Kpn*I restriction site. In parallel, large fragments containing the 3’ flanking region (FR) of δ-amastin, the pUMA1467 backbone^68^, the 5’ FR of δ-amastin, the neomycin resistance gene (*neo*), the glyceraldehyde 3-phosphate dehydrogenase intergenic region of *A. deanei* (GAPDH-IR), and eGFP were amplified from the plasmid pAEX-eGFP^28^ using forward primers containing a 5’ *Kpn*I-*Bsa*I extension and reverse primers containing a 3’ *Xho*I-*Bsa*I extension (598/599 for pAdea043 and 763/764 for pAdea235). After restriction of the fragments with *Xho*I and *Kpn*I, a total of 20-60 fmol of each of the gel-purified *lacZ*α expression cassette and the 598/599 fragment or 763/764 fragment were mixed and ligated with T4 DNA ligase to generate pAdea043 and pAdea235, respectively.

Then, each of the POI-encoding sequences were amplified from *A. deanei* gDNA with primers containing a *Bsa*I recognition site followed by 4 nucleotides complementary to the pAdea043 insertion site for the N-terminal tagging or the pAdea235 insertion site for the C-terminal tagging with eGFP (**Supplementary Table 4**). The resulting PCR fragments were extracted from agarose gels and cloned into the tagging vectors by Golden Gate ligation^69^ using equimolar or a 3:1 ratio (insert:tagging vector). In a few cases, the cloning strategy was modified, and vectors assembled by multi-fragment Golden Gate or Gibson assembly as indicated in **Supplementary Fig. 5**. *Escherichia coli* Top10 cells were transformed with the resulting vectors, transformants selected on LB agar plates containing 100 μg/ml ampicillin, and 80 μg/ml X-Gal and 0.5 mM IPTG for blue-white selection of successful ligation events if needed.

The plasmid pAdea119 containing a cassette to express ETP1 N-terminally tagged with mSCARLET from the γ-amastin locus^28^ was generated from three fragments: the mSCARLET was amplified from vector p3615 (kindly provided by Michael Feldbrügge), *etp1* was amplified from *A. deanei* gDNA using the primer set 1087/1088, and a large fragment containing the pUMA1467 backbone, 1,000 bp of the 5’- and 3’-FR of the γ-amastin gene, the hygromycin resistance gene (*hyg*), and the GAPDH-IR was amplified from the plasmid pAdea021 which targets the γ-amastin locus for insertion and expression of mCHERRY, using the primer set 1083/348 (**Supplementary Table 4**).

For generation of homozygous ETP1, ETP2, and ETP7 knock-out mutants, the plasmids pAdea148 and pAdea156 (containing replacement cassettes for ETP1), pAdea092, pAdea093, and pAdea094 (containing replacement cassettes for ETP2), and the plasmids pAdea102 and pAdea103 (containing replacement cassettes for ETP7) were constructed (**Supplementary Fig. 5**). To this end, around 1-kbp 5’ and 3’-FRs of the respective genes were amplified from *A. deanei* gDNA, *neo* was amplified from pAdea036, and *hyg* from pAdea004 (=pAdea γ-ama/Hyg^28^) using primers described in **Supplementary Table 4**. The phleomycin resistance genes (*phleo*) was synthesized by a commercial service (Integrate DNA Technologies, IDT). Vectors, carrying in the pUMA1467 backbone a replacement cassette, in which the ORF of the POI is replaced by a resistance gene, were assembled by Golden Gate ligation. The correct nucleotide sequence of expression cassettes of all plasmids generated was verified by sequencing.

### Bioinformatic analyses of the ETP amino acid sequences

Similarity searches of the *A. deanei* ETPs against the NCBI nr protein sequence database were performed using Blastp^70^. Best bidirectional blast hits were obtained by blasting the best NCBI hit back against the *A. deanei* transcriptome dataset using TBlastn built-in Bioedit v. 7.0.5.3^71^. Predictions of transmembrane regions were obtained using TMHMM v. 2.0^72^, targeting signals using TargetP 2.0^73^ and SignalP 5.0^74^. 3D structure homology was analyzed with Phyre2^46^ and disordered protein regions predicted with IUPred2A^75^. The multiple sequence alignment of OCD amino acid sequences was generated using ClustalX 2.1 and refined manually. Unambiguously alignable sequence blocks were extracted and used for phylogenetic analysis. OCD phylogeny was inferred by maximum likelihood analysis using PhyML v2.4.5^76^ with the WAG+I+G+F model of amino acid sequence evolution (determined as most suitable with ProtTest v1.4 software^77^. The robustness of branches was tested by bootstrap analysis using 100 replicates.

### Fluorescence microscopy and 3D reconstruction of fluorescence signals in *A. deanei*

50 μl of *A. deanei* cultures grown to densities between 1-8 x 10^7^ cells/ml were mixed 1:1 with PBS containing 8% paraformaldehyde (PFA), incubated for 20 min at room temperature, washed twice with PBS and then, 20 μl of the mixture was spotted on polylysine-coated glass slides. After 30 min, slides were washed 3 times with PBS followed by incubation with 10 μg/ml Hoechst 33342 in PBS for 5 min. Slides were washed 2 more times, and finally samples were mounted in 9 μl of Prolong Diamond (Thermo). Epiluminescence microscopy was carried out on an Axio imager M.1 (Zeiss, Oberkochen, Germany) coupled to a Pursuit™ 1.4 MP Monochrome CCD camera (Diagnostic Instruments, Sterling Heights, MI, USA) and a halide lamp LQ-HXP 120 (LEj, Jena, Germany) equipped with a custom set of filters for GFP (ET470/40x, T495LPXR, ET525/50m) and Rfp/mCherry (ET560/40x, T585lp, ET630/75m) (both: Chroma, Bellow Falls, VT, USA); and DAPI (447/60 BrightLine HC, HC BS 409, 387/11 BrightLine HC) (AHF Analysentechnik, Tuebingen, Germany). Images were acquired using a 100x Plan Neofluar NA 1.3 oil M27 objective (Zeiss) and processed using the Metamorph software package v. 7.7.4.0. Or an Axio Imager.A2 (Zeiss) coupled to an AxioCam MRm (Zeiss) and an Illuminator HXP 120 V (Zeiss) equipped with Filter Set 38 HE: ET470/40, BS495, ET525/50; 43 HE: ET550/25, BS570, ET605/70; 49: ET365, BS395, ET445/50. Images were acquired using an EC Plan-Neofluar 100x/1.30 Oil Ph3 M27 objective (Zeiss) and processed with Zen Blue v2.5 software. Confocal fluorescence microscopic analyses were performed on a Leica TCS SP8 STED 3X (Leica Microsystems, Wetzlar Germany) using 93x/1.3 glycerol objective equipped with a filter NF 488/561/633 with the following settings: unidirectional scan direction X, scan speed 400-1000 Hz, frame average 2, line accumulation 2 without gain. The lasers used were a diode at 405 nm and WLL at 70%. The laser line was set for Hoechst 33342 at 8.6% (405 nm), eGFP at 10% (488 nm), and mSCARLET at 7.5% (561 nm). Emission was captured by PMT between 424 and 477 nm for the Hoechst 33342 signal; and hybrid detectors were set to capture emissions between 505 and 548 nm for eGFP and 589 and 621 nm for mSCARLET. Images of both 2D and 3D representations were processed in the Leica X software v.3.5.2.18963 and deconvoluted on the Huygens Professional v. 16.10 on default confocal settings setting, except the manual mode threshold for background extraction was set based on the cytosolic background signal.

### Measurement of protein concentration, SDS-polyacrylamide gel electrophoresis (PAGE), and Western blot analysis

Protein concentrations in the samples were determined using the Pierce 660 nm Protein Assay Reagent (Thermo Fisher Scientific) in 96-well plates by the absorbance in an Infinite M200 plate reader (TECAN, Austria GmbH). For SDS-PAGE, protein samples were mixed with 4X sample buffer (final concentration 63 mM Tris-HCl, pH 6.8, 10 mM dithiothreitol, 10% glycerol, 2% SDS, and 0.0025% bromophenol blue), incubated for 5 min at 95 °C and 30 μg of protein loaded onto Bolt™ 4-12% Bis-Tris Plus precast gels (Thermo Fisher Scientific). Electrophoresis was performed at 180 V constant in 2-morpholin-4-ylethanesulfonic acid (MES)-SDS running buffer (50 mM MES, 50 mM Tris-HCl, pH 7.3, 0.1% SDS, and 1 mM ethylenediaminetetraacetic acid (EDTA). After electrophoresis, the gels were blotted onto polyvinylidene difluoride (PVDF) membranes (Amersham™ Hybond™, 0.45 nm, GE HealthCare Life Science) at 60 mA for 1 h. Membranes were blocked, incubated with a 1:1,000 dilution of mouse anti-GFP [B-2] (SantaCruz Biotechnology) or a rat anti-RFP [5F8] (Chromotek) followed by a 1:5,000 dilution of the horseradish peroxidase-conjugated secondary antibody against mouse IgG (7076, Cell Signaling Technology) or rat IgG (PA128573, Thermo Fisher Scientific), respectively, in a SNAP i.d.^®^ 2.0 (Merck-Millipore) according to the manufacturer’s instructions. Finally, membranes were covered in SuperSignal™ West Pico PLUS chemiluminescent substrate (Thermo Fisher Life Science) and chemiluminescence was detected using an ImageQuant LAS 4000 (GE Healthcare Life Science) or a ChemiDoc MP Imaging System (Bio-Rad).

## Acknowledgements

This study was supported by Deutsche Forschungsgemeinschaft grant NO 1090/1-1 (to E.C.M.N.). The authors acknowledge scientific and technical assistance of the Center for Advance Imaging (CAi) at Heinrich Heine University Düsseldorf.

## Author contributions

E.C.M.N., J.M., and G.E. designed the research. J.M., G.E., G.P., T.R., L.K., D.Z., R.W., D.K., and M.K. performed the research. J.M., G.P., and E.C.M.N. analyzed the data. E.N. and K.S. supervised the research. J.M. and E.C.M.N. wrote the manuscript.

## Data Availability Statement

The accession number for the proteome data reported in this study is: PRIDE Archive (https://www.ebi.ac.uk/pride/archive/), accession number PXD017908. Plasmids and strains generated in this study are available upon request from the authors.

