## Supplementary Material for "Evolution of a putative, host-derived endosymbiont division ring and symbiosis-induced proteome rearrangements in the trypanosomatid *Angomonas deanei*"

### Supplementary Figures

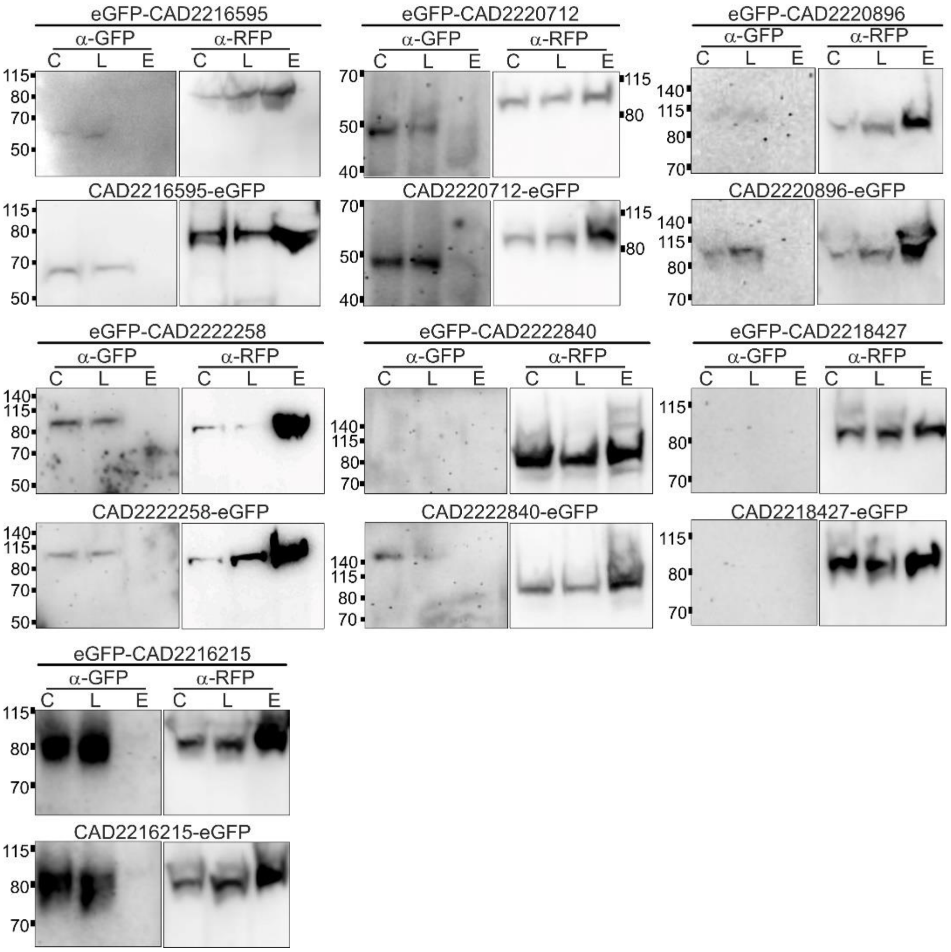


**Supplementary Figure 1: Candidate ETPs for which localization at the endosymbiont was not confirmed by Western blot.** 30 μg of protein from intact cells (C), whole cell lysate (L) or up to the Percoll step purified endosymbionts (E) were resolved by SDS-PAGE on 4-12% acrylamide gels, transferred onto PVDF-membranes, and recombinant proteins visualized using anti-GFP (α-GFP) or anti-RFP (α-RFP) antibodies with secondary antibodies conjugated to horseradish peroxidase. Note that these candidate ETPs for which localization at the endosymbiont was not confirmed did not receive an 'ETP annotation'.

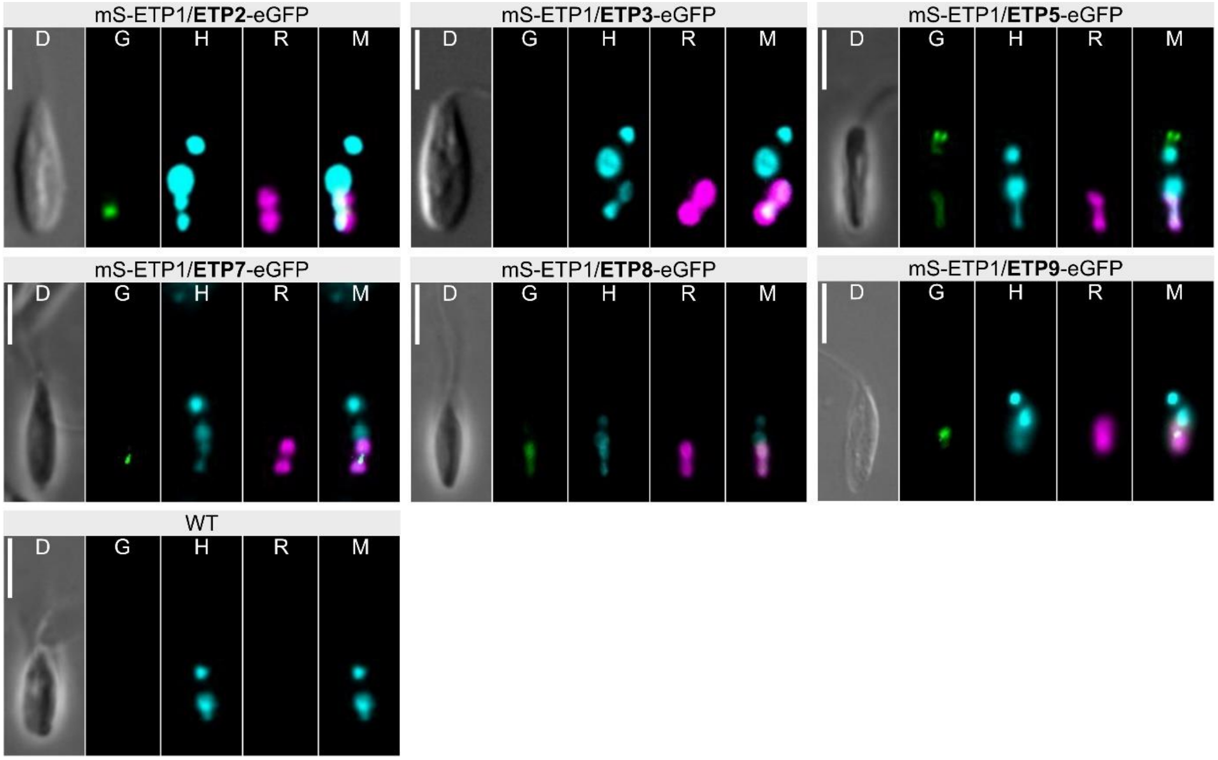


**Supplementary Figure 2: Subcellular location of the various ETPs with a C-terminal eGFP tag in *A. deanei.*** The recombinant ETPs with C-terminal eGFP tags show a green fluorescence signal corresponding to the subcellular localization observed when the eGFP tag was placed at their N-terminus. Only for ETP3-eGFP no fluorescence signal was detected. WT, wildtype; mS, mSCARLET; D, differential interference contrast; R, red channel; G, green channel; H, blue channel visualizing Hoechst 33342 staining; M, merge between the three fluorescence channels. Scale bar is 5 μm.

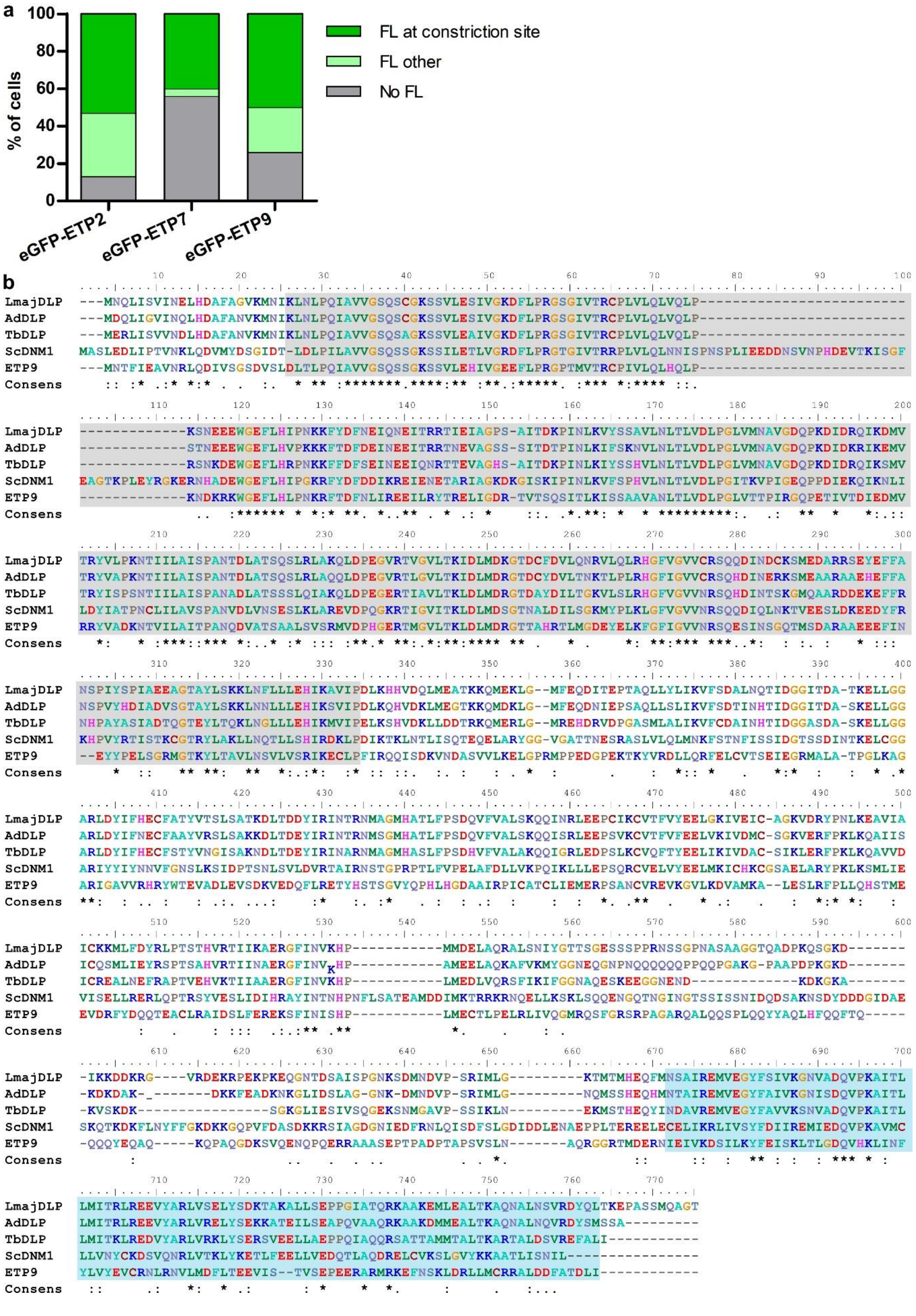
a
% of cells
0
20
40
60
80
100
FL at constriction site
FL other
No FL
eGFP-ETP2
eGFP-ETP7
eGFP-ETP9
b
LmajDLP
AdDLP
TbDLP
ScDNM1
ETP9
Consens

**Supplementary Figure 3: ETP2, ETP7, and the dynamin-like protein ETP9 form a putative division ring around the constriction site of the endosymbiont in around half of the cells in a mid-log phase culture.** (**a**) *A. deanei* cells from a mid-log phase culture expressing either eGFP-ETP2, eGFP-ETP7 or eGFP-ETP9 were fixed with 4% PFA, nucleic acid stained with Hoechst 33342 and the green fluorescence signal characterized by epifluorescence microscopy. For each cell line, 100 cells were analyzed. No FL, no fluorescence in the green channel; FL at constriction site, eGFP fluorescence signal localized at the endosymbiont constriction site; FL other, eGFP fluorescence distributed equally over the endosymbiont or restricted to one or both poles of the endosymbiont. (**b**) ClustalX amino acid sequence alignment of the dynamin-like proteins from *A. deanei* (AdDLP, CAD2218610.1 and ETP9, CAD2212698.1), *T. brucei* (TbDLP, XP_844064.1), *Leishmania major* (LmajDLP, XP_003722303.1), and *Saccharomyces cerevisiae* (ScDNM1, CAA97444.1). Consens, Clustal consensus. The N-terminal dynamin-type guanine nucleotide-binding domain and C-terminal GTPase effector domain (GED) as identified by an Expasy Prosite scan of ETP9 are shaded in grey and blue, respectively.

**CAD2220707.1 (-> ETP1)**

```
CAD2220707.1    ---------- ---------- ---------- ---------- ---------- ---------- ---------- ---------- ---------- --------MR
a814;735 K:     MTEPNAQTPP RPEQDSNNNQ TNEEEEDPRL VHPIMPAALP PPEGQIPKYL NAVYYGGRLA GIKRPVKAEL EAMGEEEAAR QKIEREERKE IKAIEKDFMR

CAD2220707.1    GVMDEERRVA KEKKEAEKKE RKEKKEAEKK RKEEEKQQQE EQQEEQTPAA EQPEAPAEEV PAQEGEEGAA TKEGGCKMCQ WLPSLHLPQL PRLPWFAKDG
a814;735 K:     GVMDEERRVA KEKKEAEKKE RKEKKEAEKK RKEEEKQQQE EQQEEQTPAA EQPEAPAEEV PAQEGEEGAA TKEGGCKMCQ WLPSLHLPQL PRLPWFAKDG

CAD2220707.1    ETPAEGEEQE EKPKVVPRRS VIGPRPNATK RPYLMIMAEE DLPPEEEVEK WRQEQRILAE EEKARQELEK ARKAEAKKAE KEAKKLAKQA KKDSKKYAKE
a814;735 K:     ETPAEGEEQE EKPKVVPRRS VIGPRPNATK RPYLMIMAEE DLPPEEEVEK WRQEQRILAE EEKARQELEK ARKAEAKKAE KEAKKLAKQA KKDSKKYAKE

CAD2220707.1    HKNDPPKEEE EEKKEEVATP QPSASPAASK AATPSPARPA YVPKAATPAA SRDASPSQPA TTAAAEPAAT PASDVVEKKD A
a814;735 K:     HKNDPPKEEE EEKKEEVATP QPSASPAASK AATPSPARPA YVPKAATPAA SRDASPSQPA TTAAAEPAAT PASDVVEKKD A
```

**CAD2221027.1 (-> ETP2)**

```
CAD2221027.1    MDYDEDEYFD NGMGFQFNNN NNNNFIPPVT PPPMYTMPAE STFPNRTNTI PVNNENSIPG QPAYTPTVAS QPGYGLGFPP SASPNKPLAE TPRERSIHSS
a8985;116 K:    MDYDEDEYFD NGMGFQFNNN NNNNFIPPVT PPPMYTMPAE STFPNRTNTI PVNNENSIPG QPAYTPTVAS QPGYGLGFPP SASPNKPLAE TPRERSIHSS

CAD2221027.1    NHGGEMRTPL RSNVGADPSI TNRFFNGPTP KRASPSVPTN NTLHTSLNMS RASSSREPVV PASMLGAPGA NPYYMFDNNN NNSMPPPPQQ PTPSPVRVAN
a8985;116 K:    NHGGEMRTPL RSNVGADPSI TNRFFNGPTP KRASPSVPTN NTLYTSLNMS RASSSREPVV PASMLGAPGA NPYYMFDNNN NNNE...... ..........

CAD2221027.1    TSDNSLNSGA RNTNNNNNEA LEQPYNVSVP PHTATVPPPP EALQVHRTPV NQSHTSSSNA HSGYVNSPSA VSSIHHPTDR LAVVTPAQNS SFASSNTAST
a8985;116 K:    .......... .......... .......... .......... .......... .......... .......... .......... .......... ..........

CAD2221027.1    TPSKFIYPRS AQRPPIILRH EVPQEDVPPA RTPFCYRRSS AAAIAV
a8985;116 K:    .......... .......... .......... .......... ......
```

**CAD2213480.1 (-> ETP3)**

```
CAD2213480.1    MIDTKFHCLI TSKERNVFDR KKSVLSVKTP SNPFSFSYNA VVSECQAWFD SWLGSVSTSS NHCVLMHAMN TESGIRLFEG TVNQIRTKLA ATGPLEIAVS
a2301;422 K:    MIDTKFHCLI TSKERNVFDR KKSVLSVKTP SNPFSFSYNA VVSECQAWFD SWLGSVSTSS NHCVLMHAMN TESGIRLFEG TVNQIRTKLA ATGPLEIAVS

CAD2213480.1    RVYSWGCMDF ISDSCIEENR HVDAGVNNLL TELSFKALES ADFHKLLKKC TSMMSGHCIV HVEIQCGNAR CSINLVPPLK DTNVILLKDL LRLRSQLLSS
a2301;422 K:    RVYSWGCMDF ISDSCIEENR HVDAGVNNLL TELSFKALES ADFHKLLKKC TSMMSGHCIV HVEIQCGNAR CSINLVPPLK DTNVILLKDL LRLRSQLLSS

CAD2213480.1    SAENSECRLF PLLIPHHDTK APCQLHVISL DDRDSSSDFH LSAWFGSSST TDDEFWQSKF RKLQTVSPAA NARKNEFKCD AQMAEKLRST RRTTDRCVSL
a2301;422 K:    SAENSECRLF PLLIPHHDTK APCQLHVISL DDRDSSSDFH LSAWFGSSST TDDEFWQSKF RKLQTVSPAA NARKNEFKCD AQMAEKLRST RRTTDRCVSL

CAD2213480.1    GAVENTDQSE QSLDPVLNAS SIEENCEQVS RTLALIKAML EEHNRREVKE HIPLSERSVP KGEDGEKRDF KVNSCADLVP LIKAKLSALE RERRKYERAV
a2301;422 K:    GAVENTDQSE QSLDPVLNAS SIEENCEQVS RTLALIKAML EEHNRREVKE HIPLSERSVP KGEDGEKRDF KVNSCADLVP LIKAKLSALE RERRKYERAV

CAD2213480.1    EDLRSAIPDS RSEVGNDDRA EDEGEECESV HSNNSLSAAG KPVEPIGVLS PSQPSLLSPD STISGSPYFD PFVSKVKKHI IKTTEEAKQM RHAIDVTFEA
a2301;422 K:    EDLRSAIPDS RSEVGNDDRA EDEGEECESV HSNNSLSAAG KPVEPIGVLS PSQPSLLSPD STISGSPYFD PFVSKVKKHI IKTTEEAKQM RHAIDVTFEA

CAD2213480.1    LGGDSSIAKA DPIIATKSLI ELAGDTSESI TQVKQLLDLE NDAVEHREKM STVLDGVREK LKKVDEVQKE ATTIVESFTK ADEEETTSVS GEGGAAVVER
a2301;422 K:    LGGDSSIAKA DPIIATKSLI ELAGDTSESI TQVKQLLDLE NDAVEHREKM STVLDGVREK LKKVDEVQKE ATTIVESFTK ADEEETTSVS GEGGAAVVER

CAD2213480.1    KHIQRKVKPM SPEEAASDAN MTLTSTLYFG VLPRSIKQNP STSLVLADIT PEAVRNFKRE MKQKMQELDE ANNALQVCVK LMKPGGKAPS THRDLVTEAL
a2301;422 K:    KHIQRKVKPM SPEEAASDAN MTLTSTLYFG VLPRSIKQNP STSLVLADIT PEAVRNFKRE MKQKMQELDE ANNALQVCVK LMKPGGKAPS THRDLVTEAL

CAD2213480.1    RLCQDANKAL TLTDQLFDDF APDDDGASPQ STITSLSGET IPKNKLTLAT RTEALVEAMK TLQNDSESCF EENDNLQRRV VFLLQQEERL QEEREAAVKR
a2301;422 K:    RLCQDANKAL TLTDQLFDDF APDDDGASPQ STITSLSGET IPKNKLTLAT RTEALVEAMK TLQNDSESCF EENDNLQRRV VFLLQQEERL QEEREAAVKR

CAD2213480.1    EQFLGNIQEK LNETEEELTT RDEELKRLQE ELKEATQKCE EATKASEMKD NIRNKRDSLL TETEYLQQDV EDLLMTKGAI EEVLGDILVS LESAVEDLEV
a2301;422 K:    EQFLGNIQEK LNETEEELTT RDEELKRLQE ELKEATQKCE EATKASEMKD NIRNKRDSLL TETEYLQQDV EDLLMTKGAI EEVLGDILVS LESAVEDLEV

CAD2213480.1    QESAMLTEVQ RAESVLRLSS TSAYAQEQKV LAKLLEAVEK QSN
a2301;422 K:    QESAMLTEVQ RAESVLRLSS TSAYAQEQKV LAKLLEAVEK QSN
```

**CAD2216595.1**

```
CAD2216595.1    MLKGASEVEL RKFQQLVNHW HDPNGALRSL HLFNPIRVGY INDMVRRYGR RVAATGDQSS GNFDYLLRSG ANGSQAGYSA FLANTSDGSK LRVLDVGCGG
a2685;461 K:    MLKGASEVEL RKFQQLVNHW HDPNGALRSL HLFNPIRVGY INDMVRRYGR RVAATGDQSS GNFDYLLRSG ANGSQAGYSA FLANTSDGSK LRVLDVGCGG

CAD2216595.1    GILSESLFRL GASVTGIDLV EESVAVANER KGMVLANLNQ SSPLYREEDL TFRQASLFEV LEEESKADGG YDVVVASEVV EHVDDARAFV KALGDVTKAR
a2685;461 K:    GILSESLFRL GASVTGIDLV EESVAVANER KGMVLANLNQ SSPLYREEDL TFRQASLFEV LEEESKADGG YDVVVASEVV EHVDDARAFV KALGDVTKAR

CAD2216595.1    GGLLIVSTME KSICSFLSHI VVAETLTGIV APRHPTTGGK FINKKDLSEY LAQHNHIAEV DHKYIASFPD PFQSAATRNL QLQFKLTNAV NTGHYLWTGL
a2685;461 K:    GGLLIVSTME KSICSFLSHI VVAETLTGIV APGTHDW-GK FINKKDLSEY LAQHNHIAEV DHKYIASFPD PFQSAATRNL QLQFKLTNAV NTGHYLWTGL

CAD2216595.1    KQ
a2685;461 K:    KQ
```

**CAD2216818.1-CAD2216821.1 (-> ETP5)**

```
CAD2216818.1    MATTLEEFSA KLDRLDQEFA KKMEEQNKKF FADKPDDSTL SPEMKEHYEK FEKMIQEHTD KFNNKMHEHS EHFKQKFAEL LEQQKNAQLP K
CAD2216819.1    MATTLEEFSA KLDRLDQEFA KKMEEQNKKF FADKPDDSTL SPEMKEHYEK FEKMIQEHTD KFNNKMHEHS EHFKQKFAEL LEQQKNAQLP K
CAD2216820.1    MATTLEEFSA KLDRLDQEFA KKMEEQNKKF FADKPDDSTL SPEMKEHYEK FEKMIQEHTD KFNNKMHEHS EHFKQKFAEL LEQQKNAQLP K
CAD2216821.1    MATTLEEFSA KLDRLDQEFA KKMEEQNKKF FADKPDDSTL SPEMKEHYEK FEKMIQEHTD KFNNKMHEHS EHFKQKFAEL LEQQKNAQLP K
a45;12605 K:    MATTLEEFSA KLDRLDQEFA KKMEEQNKKF FADKPDDSTL SPEMKEHYEK FEKMIQEHTD KFNNKMHEHS EHFKQKFAEL LEQQKNAQLP K
```

**CAD2220712.1**

```
CAD2220712.1    MLRRFSPRLC AAHHQNDHYT NNFFQKLRTR LMDRIPQTGH ETTARYKWAT HVYWYLLDPL LRCHHYYYRR KVVVDRFLER NSVFANTIFG ILLGLTVYFL
a8234;114 K:    MLRRFSPRLC AAHHQNDHYT NNFFQKLRTR LMDRIPQTGH ETTARYKWAT HVYWYLLDPL LRCHHYYYRR KVVVDRFLER NSVFANTIFG ILLGLTVYFL

CAD2220712.1    LAELVLPTAA DDEHNKNKIM HPHNMEIYSP QDNAQIVVDC IGTSTTKELP AFQLMRLKRI IMGRVLEVAD LSEVRRQREE TEKLKALLAK
a8234;114 K:    LAELVLPTAA DDEHNKNKIM HPHNMEIYSP QDNAQIVVDC IGTSTTKELP AFQLMRLKRI IMGRVLEVAD LSEVRRQREE TEKLKALLAK
```

**CAD2217314.1 (-> ETP7)**

```
CAD2217314.1    MLQSLRDKYD DHKRRVDERE RLRNDPTSDP NYYQDVINRS LLGEEVAGRP NVTPKTRRPH HDPTADPNYH QDVINRSLLN FNSETTPPPK EPKKGWFGLN
a3942;267 K:    MLQSLRDKYD DHKRRVDERE RLRNDPTSDP NYYQDVINRS LLGEEVAGRP NVTPKTRRPH HDPTADPNYH QDVINRSLLN FNSETTPPPK EPKKGWFGLN

CAD2217314.1    NVVQGALEKC KLLKEELFDD DKTDDPYDSE SNRAYRSKFA DYGDAGIPFS LCDNLNGDST YYYAKSNSCS SSRLPSSSKS KSRSTKHSSS PETVMSNNGR
a3942;267 K:    NVVQGALEKC KLLKEELFDD DKTDDPYDSE SNRAYRSKFA DYGDAGIPFS LCDNLNGDST YYYAKSNSCS SSRLPSSSKS KSRSTKHSSS PETVMSNNGR

CAD2217314.1    EYSHSRPPLD PPIKSYVGSN LNVVNNSNNT SSAFYYYHHC NRQSESNTHS SVNVDLSNIE NQVAPSVPPT APPFVLEKME SEVSTRKSYS VLPSNSDRVP
a3942;267 K:    EYSHSRPPLD PPIKSYVGSN LNVVNNSNNT SSAFYYYHHC NRQSESNTHS SVNVDLSNIE NQVAPSVPPT APPFVLEKME SEVSTRKSYS VLPSNSDRVP

CAD2217314.1    PYTIEEMHSK SRVNRVSFLD SSRSPSVAPS VVYKVMETPE ERRHLFIVKC LPHLGVVGTC ALYGNVYVET GGTFDPKQKE MHPPFHDAMG LFQMNKEMRA
a3942;267 K:    PYTIEEMHSK SRVNRVSFLD SSRSPSVAPS VVYKVMETPE ERRHLFIVKC LPHLGVVGTC ALYGNVYVET GGTFDPKQKE MHPPFHDAMG LFQMNKEMRA

CAD2217314.1    YFEKYCAEFY IRSHHLTATI VTTPSTAGGV RRDFLDPDTM DAQLTFVINE IATGSYIGAG NAATLRKSFY LLKKDISHND MRVTLTKKEY VSPPMVEAAA
a3942;267 K:    YFEKYCAEFY IRSHHLTATI VTTPSTAGGV RRDFLDPDTM DAQLTFVINE IATGSYIGAG NAATLRKSFY LLKKDISHND MRVTLTKKEY VSPPMVEAAA

CAD2217314.1    EMFCNLFERP SIPHLDRRKI SSVECLEYIR EQGLTVRM
a3942;267 K:    EMFCNLFERP SIPHLDRRKI SSVECLEYIR EQGLTVRM
```

**Supplementary Fig. 4 (continues)**

**CAD2216283.1 (-> ETP8)**

```
CAD2216283.1     ---------- ---------- ---------- ---------- ---------- ---------- ---------- ---------- ---------- ----------
RACE a6023;145   MSVVSANSNV RTVTGGNYVA QNVAVEVVKG TPFDHMIPVI TDLETKEGEP SNPSSEAKIK KLEDAGLKVQ RLPISYKGKN GDEIHEQLLP FVIHYKMQLL

CAD2216283.1     ---------- ---------- --MTSFHFGI PLKEKILLQF STLEARKLLG NTTESMNDEE TVDFSATRHL APEDLGVIND SIVNFESTVN KLLELTQTPG
RACE 6023;145    LLSERTQRKV VSNGEMTSRS NIMTSFHFGI PLKEKILLQF STLEARKLLG NTTESMNDEE TVDFSATRHL APEDLGVIND SIVNFESTVN KLLELTQTPG

CAD2216283.1     KSSK
RACE 6023;145    KSSK
```

**CAD2212698.1 (->ETP9)**

```
CAD2212698.1     MNTFIEAVNR LQDIVSGSDV SLDLTLPQIA VVGSQSSGKS SVLEHIVGEE FLPRGPTMVT RCPIVLQLHQ LPKNDKRKWG EFLHLPNKRF TDFNLIREEI
a3170;304 K:     MNTFIEAVNR LQDIVSGSDV SLDLTLPQIA VVGSQSSGKS SVLEHIVGEE FLPRGPTMVT RCPIVLQLHQ LPKNDKRKWG EFLHLPNKRF TDFNLIREEI

CAD2212698.1     LRYTRELIGD RTVTSQSITL KISSAAVANL TLVDLPGLVT TPIRGQPETI VTDIEDMVRR YVADKNTVIL AITPANQDVA TSAALSVSRM VDPHGERTMG
a3170;304 K:     LRYTRELIGD RTVTSQSITL KISSAAVANL TLVDLPGLVT TPIRGQPETI VTDIEDMVRR YVADKNTVIL AITPANQDVA TSAALSVSRM VDPHGERTMG

CAD2212698.1     VLTKLDLMDR GTTAHRTLMG DEYELKFGFI GVVNRSQESI NSGQTMSDAR AAEEEFINEY YPELSGRMGT KYLTAVLNSV LVSRIKECLP FIRQQISDKV
a3170;304 K:     VLTKLDLMDR GTTAHRTLMG DEYELKFGFI GVVNRSQESI NSGQTMSDAR AAEEEFINEY YPELSGRMGT KYLTAVLNSV LVSRIKECLP FIRQQISDKV

CAD2212698.1     NDASVVLKEL GPRMPPEDGP EKTKYVRDLL QRFELCVTSE IEGRMALATP GLKAGARIGA VVRHRYWTEV ADLEVSDKVE DQFLRETYHS TSGVYQPHLH
a3170;304 K:     NDASVVLKEL GPRMPPEDGP EKTKYVRDLL QRFELCVTSE IEGRMALATP GLKAGARIGA VVRHRYWTEV ADLEVSDKVE DQFLRETYHS TSGVYQPHLH

CAD2212698.1     GDAAIRPICA TCLIEMERPS ANCVREVKGV LKDVAMKALE SLRFPLLQHS TMEEVDRFYD QQTEACLRAI DSLFEREKSF INISHPLMEC TLPELRLIVQ
a3170;304 K:     GDAAIRPICA TCLIEMERPS ANCVREVKGV LKDVAMKALE SLRFPLLQHS TMEEVDRFYD QQTEACLRAI DSLFEREKSF INISHPLMEC TLPELRLIVQ

CAD2212698.1     GMRQSFGRSR PAGARQALQQ SPLQQYYAQL HFQQFTQQQQ YEQAQKQPAQ GDKSVQENQP QERRAAASEP TPADPTAPSV SLNAQRGGRT MDERNIEIVK
a3170;304 K:     GMRQSFSRSR PAGARQALQQ SPLQQYYAQL HFQQFTQQQQ YEQAQKQPAQ GDKSVQENQP QERRAAASEP TPADPTAPSV SLNAQRGGRT MDERNIEIVK

CAD2212698.1     DSILKYFEIS KLTLGDQVHK LINFYLVYEV CRNLRNVLMD FLTEEVISTV SEPEERARMR KEFNSKLDRL LMCRRALDDF ATDLI
a3170;304 K:     DSILKYFEIS KLTLGDQVHK LINFYLVYEV CRNLRNVLMD FLTEEVISTV SEPEERARMR KEFNSKLDRL LMCRRALDDF ATDLI
```

**CAD2220896.1**

```
CAD2220896.1     ---------- ---------- ---------- ---------- ---------- ---------- ---------- ---------- ---------- ----------
a5402_199 K:     MRSFARRSLV PCLASMTHGL QRPSPSWCSG RSCQSAPPPQ EEDSHHHHVE ERPRQFQQDM ELRQLVQSAV PLQSTEDIKT MLHGVKNAKE VRHILETVVY

CAD2220896.1     ---------- ---------- ---------- ---------- ---------- ---------- ---------- ---------- ---------- ----------
a5402_199 K:     RHHLCGDVLN GHGFRVTIVQ SLTLVAPRVQ AMEEWFDCLG RFRKLNFLLT RTFAAEGLHL IKEWLTHRFN TEGRTPFLLT TGTAYIRELM YWCHEDKLVF

CAD2220896.1     ---------- ---------- ---------- ---------- ---------- ---------- ---MEEVLDL LRNDIPSRPN FNLLFRLMDY YFATDNVEKL
a5402_199 K:     DHVLYTRIVF LLTIIVSFFD RQNLYRTSFP SDFVKRDGIV TEWIHTNERC VDYDECVVQC DALMEEVLDL LRNDIPSRPN FNLLFRLMDY YFATDNVEKL

CAD2220896.1     IAVMEDAHEY GVTVAESSTA KLMQLACAFN YPTAPELFLR WRVSLPQCAI ASPDISRLLF YYSRSGGGLP CPACGEKYNH RNVNVYHWMA TPPHQRQCPV
a5402_199 K:     IAVMEDAHEY GVTVAESSTA KLMQLACAFN YPTAPELFLR WRVSLPQCAI ASPDISRLLF YYSRSGGGLP CPACGEKYNH RNVNVYHWMA TPPHQRQCPV

CAD2220896.1     LHMARNRKGD LEADPALPQN RDWSERALQL RELSTARSIE WGTQEWRGFL GCFMFAPTEK AMQAKALLDQ SMGAAQMDDF LRAAYIRLLR YHAPELSLPV
a5402_199 K:     LHMARNRKGD LEADPALPQN RDWSERALQL RELSTARSIE WGTQEWRGFL GCFMFAPTEK AMQAKALLDQ SMGAAQMDDF LRAAYIRLLR YHAPELSLPV

CAD2220896.1     LRQWEDSGIR MSPIVLQEAL MAAVTLDDSV QRLESLTHTW DLLREKGSYV MPFTRRYVER RRDALQQQAP LSGEESHIIR EVVEMRPRTV SLLDRKDSSS
a5402_199 K:     LRQWEDSGIR MSPIVLQEAL MAAVTLDDSV QRLESLTHTW DLLREKGSYV MPFTRRYVER RRDALQQQAP LSEEESHIIR EVVEMRPRTV SLLDRKDSSS

CAD2220896.1     DFVIGTSKKN IYIPKRSLKE SERRRDQRRQ RSQSE
a5402_199 K:     DFVIGTSKKN IYIPKRSLKE SERRRDQRRQ RSQSE
```

**CAD2222258.1**

```
CAD2222258.1     ---------- ---------- ---------- ---------- ---------- ---------- ---------- ---------- ---------- ----------
a5695_215 K:     MTDSDQHLSN ETQLEFTNND SNDQNPATDV ADMNLSEFRR VVQESQPQDN HLPAPPRAVL KYIPEVTDDF IRNFLLRNNM LKTLEQFEVE WYSRFGSAAS

CAD2222258.1     ---------- ---------- ---------- ---------- ---------- ---------- ---------- ---------- -----MKRRC DGLLKQKTVL
a5695_215 K:     RDVPLVPDNY LETAQLSHRI ELLERDLREH AELTTTLNKQ WRQAKKERDF HKANHSRVVQ EKNLKTKVLK QTDRNAADLT PTLEEMKRRC DGLLKQKTVL

CAD2222258.1     SLEKDKLAAQ VEQLEKRLSE AERRLEGEEG ESSTTSGKPK RSKSSQKKEN TSSSAARKRG VAAAVSAART GTSTTKTADG STTDGFVWPP DERPNTEAAP
a5695_215 K:     SLEKDKLAAQ VEQLEKRLSE AERRLEGEEG ESSTTSGKPK RSKSSQKKEN TSSSAARKRG VAAAVSAART GTSTTKTADG STTDGFVWPP DERPNTEAAP

CAD2222258.1     PSLPSHVTQW SNTHTFTAHA MSVTKIALHP RKPAVASCSD DGTWRLSTVP EGELILSGEG HQNWVAAVAM HPAGTMVATG SGDKTVKLWD FATNSCANTL
a5695_215 K:     PSLPSHVTQW SNTHTFTAHA MSVTKIALHP RKPAVASCSD DGTWRLSTVP EGELILSGEG HQNWVAAVAM HPAGTMVATG SGDKTVKLWD FATNSCANTL

CAD2222258.1     RSHTDGVWSV DFQETGALLA SGSLDTTARV WDVEMGKCRQ TLRGHVEAVN AVQWCVGTNI LCTGSGDKTV SLWDTRMNCC ATTLYGHRSP VLSVSTLPGN
a5695_215 K:     RSHTDGVWSV DFQETGALLA SGSLDTTARV WDVEMGKCRQ TLRGHVEAVN AVQWCVGTNI LCTGSGDKTV SLWDTRMNCC ATTLYGHRSP VLSVSTLPGN

CAD2222258.1     QVLASCDTEG TVVVWDIRKL EQKQSFACGP SPANCVTFDG VGEWLAVGSD DSILRLIDLE KETVSELRGH EDGVLCCAFD PAMRFLVSSG SDCTVRYWN
a5695_215 K:     QVLASCDTEG TVVVWDIRKL EQKQSFACGP SPANCVTFDG VGEWLAVGSD DSILRLIDLE KETVSELRGH EDGVLCCAFD PAMRFLVSSG SDCTVRYWN
```

**CAD2222840.1**

```
CAD2222840.1     MTILENPAPS SSFRNWQNIA TGYGAFCVQE RTAAARRPSV ISFAPAEKPF RIIFSHCPID FLSLTEKLSA KSSTTGLPSS AGSGNKRFVV AVASSLATIL
a5888_177 K:     MTILENPAPS SSFRNWQNIA TGYGAFCVQE RTAAARRPSV ISFAPAEKPF RIIFSHCPID FLSLTEKLSA KSSTTGLPSS AGSGNKRFVV AVASSLATIL

CAD2222840.1     EDWGWVMKGW QTVGQRTDSA MKRRTRTSRT AGSSDDALTR EEWSEVVQVI KEQLEVEREA EYVETTTENM LDEEDGGKKP LSPSASVRDA ALLVSRDRRL
a5888_177 K:     EDWGWVMKGW QTVGQRTDSA MKRRTRTSRT AGSSDDALTR EEWSEVVQVI KEQLEVEREA EYVETTTENM LDEEDGGKKP LSPSASVRDA ALLVSRDRRL

CAD2222840.1     SVDDSLDTTP EVSGAPTRDD TFLAELEEEP ANKVTGVYFY TSDIFDIILL RHIDVWAWSS PHVTILMIMV CFAFPWNQAL VILLAEALPL SLLQRRRAVD
a5888_177 K:     SVDDSLDTTP EVSGAPTRDD TFLAELEEEP ANKVTGVYFY TSDIFDIILL RHIDVWAWSS PHVTILMIMV CFAFPWNQAL VILLAEALPL SLLQRRRAVD

CAD2222840.1     KRTKKRDQRH LETFSKLFSK LGHLKEEEPE TKVNPFKQES DVSVGEEEEE PVKPEDPLVT RVGNFMGTFL AKVLFVAVLL FALFNLILPR SVDLLAVADA
a5888_177 K:     KRTKKRDQRH LETFSKLFSK LGHLKEEEPE TKVNPFKQES DVSVGEEEEE PVKPEDPLVT RVGNFMGTFL AKVLFVAVLL FALFNLILPR SVDLLAVADA

CAD2222840.1     VVGKVVGAAL LLCGVRQVLK PPVVNRRPLS RKAITNLLAS TQRNIHPHID NPSHRYVRPE LSESEKSMVD QLRNFGAPYP LILLRPLRDL SVATGWTPVV
a5888_177 K:     VVGKVVGAAL LLCGVRQVLK PPVVNRRPLS RKAITNLLAS TQRNIHPHID NPSHRYVRPE LSESEKSMVD QLRNFGAPYP LILLRPLRDL SVATGWTPVV

CAD2222840.1     LGEKVRCEHR YSNNDTVHSI RYSVNVPLAN VGTMQAVLLD DVDGQFDELH SSNLYTWDTL LQSRQILKKL DYNLYVVRYQ RRSSMWGVPS SDLDQFVIPA
a5888_177 K:     LGEKVRCEHR YSNNDTVHSI RYSVNVPLAN VGTMQAVLLD DVDGQFDELH SSNLYTWDTL LQSRQILKKL DYNLYVVRYQ RRSSMWGVPS SDLDQFVIPA

CAD2222840.1     VVLNSEQQKA LNISDITPRP PAAEGRRKGH GNLRAFLQCA IPCPESFSAP YVEEAAEGKG RSPTSKEEPT KIAATSLLLG LEEADGSLTL CLYRSYSGVT
a5888_177 K:     VVLNSEQQKA LNISDITPRP PAAEGRRKGH GNLRAFLQCA IPCPESFSAP YVEEAAEGKG RSPTSKEEPT KIAATSLLLG LEEADGSLTL CLYRSYSGVT

CAD2222840.1     RSKFLEEQRL IMGEASTHCL AYLLEATGYQ PSQIADFSNA SGFQPFFCRQ PFTDLPAEEE ENVDPGDDTL ELPAMGARNG VSPERDQGDG MIKRQRSLRF
a5888_177 K:     RSKFLEEQRL IMGEASTHCL AYLLEATGYQ PSQIADFSNA SGFQPFFCRQ PFTDLPAEEE ENVDPGDDTL ELPAMGARNG VSPERDQGDG MIKRQRSLRF

CAD2222840.1     MSTLPLPTVC QTVVRDALQS NLWRLRKTRG GVRVMECTRT VPGIWDGTPH VSVFCAQVVV SCNFFHVMRT LSRNSAVTVY NDAVESRVPL GSQIPVAELL
a5888_177 K:     MSTLPLPTVC QTVVRDALQS NLWRLRKTRG GVRVMECTRT VPGIWDGTPH VSVFCAQVVV SCNFFHVMRT LSRNSAVTVY NDAVESRVPL GSQIPVAELL

CAD2222840.1     ATVEEESSDE EEAALTGVTS VNVSPPVSTV LKQVGEVRGT GSGVYHTAFR GKFGAVARDA ITKEHGPYFF TAKSLREHLF DATQSAEMDL LRELPNDTRM
a5888_177 K:     ATVEEESSDE EEAALTGVTS VNVSPPVSTV LKQVGEVRGT GSGVYHTAFR GKFGAVARDA ITKEHGPYFF TAKSLREHLF DATQSAEMDL LRELPNDTRM

CAD2222840.1     CVRVEEDPTL DDGETVQAGK TKNTTKKPKK KREIVPPLMD YERCHVHRRG VLCYEIPSKK EGVPQTMVMH YSCAEPSGWL PKLYANVIEV EQLLLSAEKF
a5888_177 K:     CVRVEEDPTL DDGETVQAGK TKNTTKKPKK KREIVPPLMD YERCHVHRRG VLCYEIPSKK EGVPQTMVMH YSCAEPSGWL PKLYANVIEV EQLLLSAEKF

CAD2222840.1     KTLVEYSNKE K
a5888_177 K:     KTLVEYSNKE K
```

**Supplementary Figure 4 (continues)**

**CAD2218427.1**

```
CAD2218427.1    MLKNESKPPV EDSSKIGGGS VIVEVKDTSS TTHYEGDGKA GNEPYQAEKS TGEFSYTSDT QDTPFGVSLS LLKRNMYLFI WLKAIGSYDS GAFSAVLAVE
a4999;217 K:    MLKNESKPPV EDSSKIGGGS VIVEVKDTSS TTHYEGDGKA GNEPYQAEKS TGEFSYTSDT QDTPFGVSLS LLKRNMYLFI WLKAIGSYDS GAFSAVLAVE

CAD2218427.1    NGMSDSLGLS TLDKGNLAAS VFLGNIIGCP VAGHLFGSYN EQNVLNASLI AHTVATFLFA FFPGYYYCVF FRFFIGFTLA FIVVYTPVWV DHFAPRDKKS
a4999;217 K:    NGMSDSLGLS TLDKGNLAAS VFLGNIIGCP VAGHLFGSYN EQNVLNASLI AHTVATFLFA FFPGYYYCVF FRFFIGFTLA FIVVYTPVWV DHFAPRDKKS

CAD2218427.1    IWMASHNAGV PLGIMFGYLV GAYFPSFTAI PWEWAFYLKC LLMVPTIVYV GSANPRSLNA RGPGALMTED NNSDTSSPKT RQTNLAGTTI PGDKGVLAQL
a4999;217 K:    IWMASHNAGV PLGIMFGYLV GAYFPSFTAI PWEWAFYLKC LLMVPTIVYV GSANPRSLNA RGPGALMTED NNSDTSSPKT RQTNLAGTTI PGDKGVLAQL

CAD2218427.1    KTSATTLFRR FSPLIANPVF MCAVFAMSAM YLVATGLQNF VTEYLKEEPF NASILTIMLG FGRRW..... .......... .......... ..........
a4999;217 K:    KTSATTLFRR FSPLIANPVF MCAVFAMSAM YLVATGLQNF VTEYLKEEPF NASILTIMLG FCAAVVTSPV LGVIVGGVLL DRLGGYHGNL LVASAFATAW

CAD2218427.1    .......... .......... .......... .......... .......... .......... .......... .......... .......... ..........
a4999;217 K:    GFAATIFSII CIFVTSTGWF LIVMSCVLFC GGAIIPPGAG ITMSTLPAPL RSAGAAFAQT MYNLLGNYSG PLLCGFIAKQ TGHLKYGIYT LFLCSLLGVV

CAD2218427.1    .......... .......... .......... .......... .......... .......... .......... .......... .......... ..........
a4999;217 K:    PMSFIVLIAW RRKQSGVTAD TVVVMDTIEE EGGGGVDQEM AHVPPAGGSF SVRKENKEEV LPFSRTTQKV SPLESWRRGQ ERPARLASPP GSVAVEPSAA

CAD2218427.1    .......... .......... .......... .......... .......... .......... .......... .......... .......... ...
a4999;217 K:    PTPLHTPKDS GLRTRESFLG MLDGSQEKSI PNQHAFGMDL VYSWLTAQEE TERRRTASVA GRTNLHPLQA DSVEMSPSVD SESRRRKNHN SRE
```

**CAD2216215.1**

```
CAD2216215.1    ---------- ---------- ---------- ---------- ---------- ---------- ---------- ---------- ---------- ----------
a4187_164 K:    MFSSPFNFLL HFSGAHKGRI LFTALLMSAA TFIFLLFPFF GEELVTFTTE TEKTTETPKD DFYTHCRQKL TEWEMVLFSP LIYAGAALFV ESISPDTKLL

CAD2216215.1    ---------- ---------- ---------- ---------- ---------- ---MMLDGSV ITGIVTSNAK LCGNSVARFL TDVLYLCLSV IGLAGMLIFL
a4187_164 K:    LSHAFALGAS IISYHLVSLL AHLSAYRVSM RVQHSIITSS LATILSTEFP ERVMMLDGSV ITGIVTSNAK LCGNSVARFL TDVLYLCLSV IGLAGMLIFL

CAD2216215.1    SCRLTLIIGV LIVSVQLFYL FAGKSNRKKG TEVNHAETTA HSYLTNAIRR SETISIFNCA PFVVDRLEST FADLQRLSSS LNFSIHGYAA LSSGSTNLIF
a4187_164 K:    SCRLTLIIGV LIVSVQLFYL FAGKSNRKKG TEVNHAETTA HSYLTNAIRR SETISIFNCA PFVVDRLEST FADLQRLSSS LNFSIHGYAA LSSGSTNLIF

CAD2216215.1    VIVLVLANYY HRQGSLELTD IAMYFMLFQS FVRTLSYLAE EANSFRSMIN GTAVLYQLMN WYQEVRVPAE GDDKRVCFIP EEDTTTPVEL KAVAFAYPEI
a4187_164 K:    VIVLVLANYY HRQGSLELTD IAMYFMLFQS FVRTLSYLAE EANSFRSMIN GTAVLYQLMN WYQEVRVPAE GDDKRVCFIP EEDTTTPVEL KAVAFSYPEI

CAD2216215.1    PSFLETQFQS LSTVGGEELK TKKGITDISF SVPQHSITVL FGQSGCGKST SLRILGGILQ PSGGTVNTRK NAILLEQQHA IFFGSVAENI MLKDLSLCAD
a4187_164 K:    PSFLETQFQS LSTVGGEELK TKKGITDISF SVPQQSITVL FGQSGCGKST SLRILGGILQ PSGGTVNTRK NAILLEQQHA IFFGSVAENI MLKDLSLCAD

CAD2216215.1    EHTERVQKAC AKGGCDVFIK NPFREMISNT DKPPFSGGQL QRICLARLFA NCDDCSLILF DEPTTGLDAT RVKSLLETIS LLRDTYKKTV VIASHDERVL
a4187_164 K:    ENTERVQKAC AKGGCDVFIK NPFREMISNT DKPPFSGGQL QRICLARLFA NCDDCSLILF DEPTTGLDAT RVKSLLETIS LLRDTYKKTV VIASHDERVL

CAD2216215.1    NFADHVVTFK
a4187_164 K:    NFADHVVTFK
```

**Supplementary Figure 4: Alignments of candidate ETP gene models in genome assembly GCA_903995115.1 with corresponding 5’ full-length *A. deanei* transcripts.** N-terminal extensions of ORFs deduced from transcript sequences are highlighted in blue. The transcript sequence of ETP2 is incomplete at the C-terminal end.

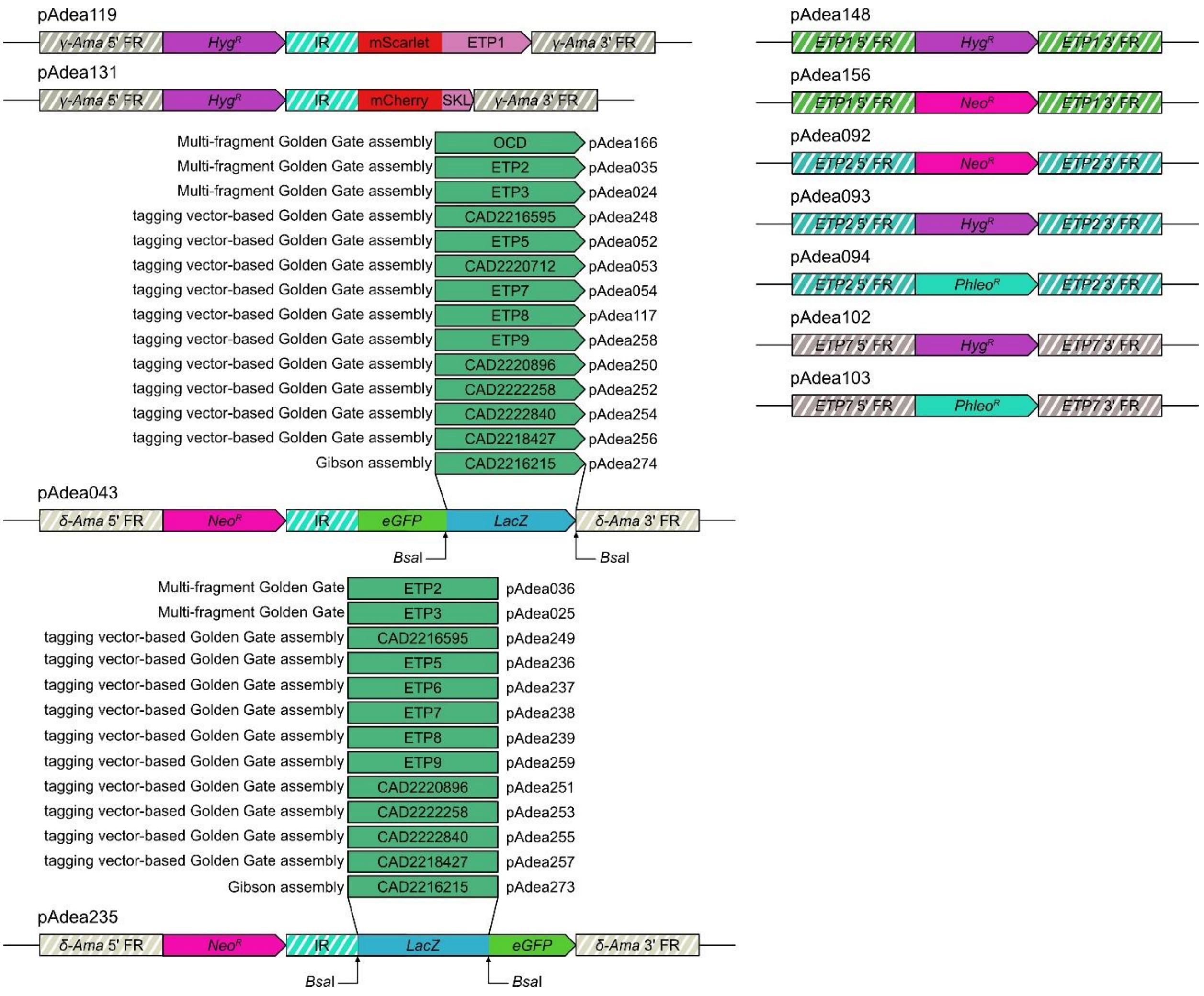


**Supplementary Fig. 5: Cassettes used for the expression of recombinant proteins and targeted gene knock-outs in *A. deanei*.** The plasmid pAdea119 contains a cassette for the expression of ETP1 (pink arrow) fused to the C-terminus of mSCARLET (red rectangle) from the $\gamma$-amastin locus; the $\gamma$-amastin flanking regions (FRs) used for homologous recombination (HR) are represented by hatched grey bars. The plasmids pAdea043 and pAdea235 contain cassettes for expression from the $\delta$-amastin locus; FRs used for HR are represented by hatched brown bars. By replacement of the *lacZ*$\alpha$ expression cassette with the POI (dark green rectangles) at the *Bsa*I sites, the POI is scarlessly fused to the C-terminus (pAdea043) or N-terminus (pAdea235) of eGFP (green rectangle/arrow). The absence/presence of the *lacZ*$\alpha$ cassette allows for blue/white selection of the resulting plasmids. Hygromycin resistance gene (*hyg*, violet arrow); neomycin resistance gene (*neo*, fuchsia arrow); the intergenic region between the glyceraldehyde 3-phosphate dehydrogenase I and II from *A. deanei* (IR, hatched cyan bar); *lacZ*$\alpha$ expression cassette (*lacZ*, blue-green rectangle). The plasmids pAdea148, pAdea156, pAdea092, pAdea093, pAdea094, pAdea102, and pAdea103 contain the resistance marker genes *hyg, neo* or *phleo* flanked by ~1,000 bp of the 5' and 3' FR of ETP1, ETP2, and ETP7 as indicated. These cassettes were used to target both alleles of the respective genes for deletion by HR.

### Supplementary Movies

**Supplementary movies 1-6. Three-dimensional reconstruction of the fluorescence signals obtained from *A. deanei* cell lines co-expressing mS-ETP1 and one of each of the remaining ETPs fused to the C-terminus of eGFP.** The signals are split into four separate channels showing the mS-ETP1 (magenta); the eGFP-POI (green); the DNA of the nucleus, kinetoplast, and endosymbiont stained with Hoechst 33342 (cyan); and the overlay of the 3 channels. Each clip shows the rotation of 360 degrees of the cell along the horizontal or vertical axis of the screen.

- MoralesEhret_SupplMovie1(ETP2_series).mp4
- MoralesEhret_SupplMovie2(ETP3_series).mp4
- MoralesEhret_SupplMovie3(ETP5_series).mp4
- MoralesEhret_SupplMovie4(ETP7_series).mp4
- MoralesEhret_SupplMovie5(ETP8_series).mp4
- MoralesEhret_SupplMovie6(ETP9_series).mp4

### Supplementary Tables

**Supplementary Table S1:** see separate Excel file MoralesEhret_SupplTable1.xlsx.

**Supplementary Table S2: Mass spectrometric identification of candidate ETPs.**

| **Accession**[a] | **Annotation** | **Exp. 1**[b] | **Exp. 2**[c] | **Chr**[d] | **N-ext [aa]**[e] | **M.W. [kDa]**[f] | **ETP**[g] |
|---|---|---|---|---|---|---|---|
| CAD2220707.1 | Hypothetical protein, cons. | 3.4/2.4 | 4.1/8.2 | 18 | 98 | 42.7 | ETP1 |
| CAD2221027.1 | Hypothetical protein, cons. | 8.2/3.5 | 6.9/8.8 | 19 | - | 53.4 | ETP2 |
| CAD2213480.1 | Hypothetical protein, cons. | 1.5/2.9 | 5.2/6.0 | 02 | - | 105.1 | ETP3 |
| CAD2216595.1 | Nodulation protein S (NodS)/Methyltransferase domain/Ribosomal protein L11 methyltransferase (PrmA), put. | 5.3/1.8 | 0.1/0.0 | 07 | - | 32.9 | |
| CAD2216818.1; CAD2216819.1; CAD2216820.1; CAD2216821.1 | Kinetoplastid membrane protein 11, put. | 0.4/0.3 | 2.7/3.4 | 07 07 07 07 | - | 11.0 | ETP5 |
| CAD2220712.1 | Hypothetical protein, cons. | 4.1/3.8 | nd | 18 | - | 22.5 | |
| CAD2217314.1 | Phage tail lysozyme, put. | 5.1/2.1 | 3.4/3.4 | 08 | - | 61.0 | ETP7 |
| CAD2216283.1 | Hypothetical protein, cons. | 5.1/2.5 | 4.5/5.9 | 06 | 122 | 22.6 | ETP8 |
| CAD2212698.1 | Dynamin family/Dynamin central region/Dynamin GTPase effector domain containing protein, putative | 1.6/1.7 | 0.5/0.4 | 01 | - | 77.4 | ETP9 |
| CAD2220896.1 | Hypothetical protein, cons. | 8.8/2.5 | nd | 19 | 263 | 73.9 | - |
| CAD2222258.1 | WD domain, G-beta repeat, put. | 3.5/2.1 | nd | 25 | 185 | 66.2 | - |
| CAD2222840.1 | Hypothetical protein, cons. | nd | 2.5/3.9 | 28 | - | 123.8 | - |
| CAD2218427.1 | Sugar (and other) transporter/Major Facilitator Superfamily/Uncharacterised MFS-type transporter YbfB/Organic Anion Transporter Polypeptide (OATP) family, putative | 7.1/3.5 | nd | 11 | - | 74.8 | - |
| CAD2216215.1 | ABC transporter transmembrane region/ABC transporter, putative | nd | 4.3/4.2 | 06 | 153 | 67.6 | - |

[a] GenBank accession number.
[b,c] Enrichment of protein in ES fractions in LC-MS/MS Experiment 1 and 2, indicated by (difference ES-WC/-log T-test p-value) as in **Fig. 1c-d**. Nd, not detected.
[d] Chromosomal localization in *A. deanei* nuclear genome assembly GCA_903995115.1.
[e] N-terminal extension of the ORF as predicted by full-length transcript by indicated number of amino acids (see **Supplementary Fig. 4**).
[f] Estimated molecular weight of the protein encoded by the full length ORF (Expasy Protparam).
[g] Newly assigned name.

**Supplementary Table S3: Primers used in this study.** For each primer, the table lists an internal primer number, its nucleotide sequence, a short description of the fragment it amplifies during PCR, and the identifier (pAdea###) of the resulting plasmid (see **Supplementary Fig. 4**).

| Number | Sequence | Fragment | pAdea |
|---|---|---|---|
| 1083 | GGTCTCAGTAATAAAAAACATACAAAACAAAAC | 3'-γ-ama FR/ pUMA1467 backbone/5'-γ-ama FR/*hyg*/GAPDH IR/ | |
| 348 | GGTCTCCTTGGATAACTGTGTTTTTTGATGAA | | |
| 1085 | GGTCTCACCAAATGGTGAGCAAGGGCGAGGC | mScarlet | 119 |
| 1086 | GGTCTCACCGTCTTGTACAGCTCGTCCATGC | | |
| 1087 | GGTCTCTACGGAACCCAACGCCCAAAC | ETP1 “allele Q-“ | |
| 1088 | GGTCTCGTTACGCATCCTTCTTCTCTA | | |
| 598 | CACGGTACCGGTCTCTAAGAGGGGGGAGAGAGACGTG | BsaI/3'-δ-ama FR/pUMA1467 backbone/5'-δ-ama FR/*neo*/GAPDH IR/eGFP/BsaI | |
| 599 | CACCTCGAGGGTCTCGCTTGTACAGCTCGTCCATGC | | 043 |
| 596 | CACCTCGAGTTTACACTTTATGCTTCCGG | XhoI/LacZ alpha with promoter/KpnI | |
| 597 | CACGGTACCTTAATGCGCCGCTACAG | | |
| 763 | CACGGTACCGGTCTCTGTGAGCAAGGGCGAGGAG | BsaI/eGFP/3'-δ-ama FR/pUMA1467 backbone/5'-δ-ama FR/*neo*/GAPDH IR/BsaI | |
| 764 | CACCTCGAGGGTCTCGTTGGATAACTGTGTTTTTTGATGAAAG | | 235 |
| 596 | CACCTCGAGTTTACACTTTATGCTTCCGG | XhoI/LacZ alpha with promoter/KpnI | |
| 597 | CACGGTACCTTAATGCGCCGCTACAG | | |
| 121 | GGTCTCGCCTGCAGTGCGTCGCCCGGCTA | 5'-δ-ama FR/*neo*/GAPDH IR/eGFP | |
| 362 | GGTCTCGTCCTTGTACAGCTCGTCCAT | | |
| 363 | GGTCTCGAGGACTACGACGAGGATGAAT | ETP2 | 035 |
| 364 | GGTCTCGCTTTTAGTTCGCCGAAAGCAC | | |
| 365 | GGTCTCCAAAGAGGGGGGAGAGAGACGT | 3'-δ-ama FR | |
| 126 | GGTCTCGCTGCGTCGCGGGGGCTGTCGCA | | |
| 121 | GGTCTCGCCTGCAGTGCGTCGCCCGGCTA | 5'-δ-ama FR/*neo*/GAPDH IR/eGFP | |
| 354 | GGTCTCCCTTGTACAGCTCGTCCATG | | |
| 355 | GGTCTCGCAAGATCGACACGAAATTTCATTGT | ETP3 | 024 |
| 356 | GGTCTCGTTCTAGTTGGACTGCTTTTCTAC | | |
| 357 | GGTCTCCAGAAGAGGGGGGAGAGAGACG | 3'-δ-ama FR | |
| 126 | GGTCTCGCTGCGTCGCGGGGGCTGTCGCA | | |
| 612 | GGTCTCGCAAGATGGCCACCACCCTTGAGGAA | ETP5 | 052 |
| 613 | GGTCTCGTCTTTTACTTGGGGAGCTGGGCGT | | |
| 614 | GGTCTCGCAAGATGCTCCGTCGATTTTCCCCA | CAD2220712 | 053 |
| 615 | GGTCTCGTCTTCTACTTGGCCAGCAGTGCCT | | |
| 616 | GGTCTCGCAAGATGCTGCAATCCTTACGGGAC | ETP7 | 054 |
| 617 | GGTCTCGTCTTTCACATGCGTACCGTTAACC | | |
| 1280 | GGTCTCGCAAGAGTGTAGTGTCCGCCAACTCCAAC | ETP8 | 117 |
| 1281 | GGTCTCGTCTTTTACTTGGACGACTTTCCAGGAGT | | |
| 121 | GGTCTCGCCTGCAGTGCGTCGCCCGGCTA | 5'-δ-ama FR/*neo*/GAPDH IR | 036 |

| | | | |
|---|---|---|---|
| 303 | GGTCTCGTTTGGATAACTGTGTTTTTTGATGA | | |
| 263 | GGTCTCGCAAATGGACTACGACGAGGAT | ETP2 | |
| 264 | GGTCTCGGTTCGCCGAAAGCACAAA | | |
| 265 | GGTCTCCGAACGTGAGCAAGGGCGAGGAG | eGFP/3'-δ-ama FR | |
| 126 | GGTCTCGCTGCGTCGCGGGGGCTGTCGCA | | |
| 121 | GGTCTCGCCTGCAGTGCGTCGCCCGGCTA | 5'-δ-ama FR/*neo*/GAPDH IR | |
| 273 | GGTCTCCTCATTTGGATAACTGTGTTTTTTGATGAAAGAAGAC | | |
| 274 | GGTCTCGATGATCGACACGAAATTTCAT | ETP3 | 025 |
| 275 | GGTCTCGGTTGGACTGCTTTTCTACTG | | |
| 276 | GGTCTCCCAACGTGAGCAAGGGCGAGGAG | EGFP/3'-δ-ama FR | |
| 126 | GGTCTCGCTGCGTCGCGGGGGCTGTCGCA | | |
| 630 | GGTCTCGCCAAATGGCCACCACCCTTGAGGAA | ETP5 | 236 |
| 893 | GGTCTCGTCACCTTGGGGAGCTGGGCGT | | |
| 632 | GGTCTCGCCAAATGCTCCGTCGATTTTCCCCA | CAD2220712 | 237 |
| 894 | GGTCTCGTCACCTTGGCCAGCAGTGCCT | | |
| 634 | GGTCTCGCCAAATGCTGCAATCCTTACGGGAC | ETP7 | 238 |
| 895 | GGTCTCGTCACCATGCGTACCGTTAACC | | |
| 1278 | GGTCTCGCCAAATGAGTGTAGTGTCCGCCAACTCC | ETP8 | 266 |
| 1279 | GGTCTCTTCACCTTGGACGACTTTCCAGGAGTCTG | | |
| 863 | GGTCTCGTGTGACTGTCAGAAAGCGGC | 5' FR scaffold 1854 | |
| 844 | GGTCTCCTTAGAAGAACTCGTCAAGAAGG | | |
| 524 | GGTCTCCCTAAGGAAAAGATAAATAGACGTTAAAAAAG | *neo*/3' FR scaffold 1854 | 165 |
| 595 | GGTCTCCGCACAAATAATGGTTTTGAGGCT | | |
| 590 | GGTCTCGGTGCAGGTCTAGATATCGGATC | pUMA 1467 | |
| 862 | GGTCTCGCACAGGTGAGCTCGAATTCA | | |
| 1835 | ATGGTAGGTCTCACAAGATGCTGAAAGGCGCCTCTGAGGTGG | CAD2216595 | 248 |
| 1836 | ATGGTAGGTCTCATCTTTTACTGTTTCAGTCCAGTCCACAGA | | |
| 1837 | ATGGTAGGTCTCACCAAATGCTGAAAGGCGCCTCTGAGGTGG | CAD2216595 | 249 |
| 1838 | AGTGTAGGTCTCATCACCTGTTTCAGTCCAGTCCACAGATAG | | |
| 1849 | GGTCTCACAAGATGAGAAGTTTCGCTAGAAGGTC | CAD2220896 | 250 |
| 1850 | GGTCTCAtcttTTACTCACTTTGTGATCGTTGGC | | |
| 1851 | GGTCTCACCAAATGAGAAGTTTCGCTAGAAGGTC | CAD2220896 | 251 |
| 1852 | GGTCTCATCACCTCACTTTGTGATCGTTGG | | |
| 1853 | GGTCTCACAAGATGACAGACAGCGATCAACAC | CAD2222258 | 252 |

| | | | |
|---|---|---|---|
| 1854 | GGTCTCAtcttCTAATTCCAGTAGCGCACCGTA | | |
| 1855 | GGTCTCACCAAATGACAGACAGCGATCAACAC | CAD2222258 | 253 |
| 1856 | GGTCTCATCACATTCCAGTAGCGCACCGTAC | | |
| 1857 | GGTCTCACAAGATGACGATCTTGGAGAATCCAGC | CAD2222840 | 254 |
| 1858 | GGTCTCAtcttTTACTTTTCTTTATTCGAATACTCCACG | | |
| 1859 | GGTCTCACCAAATGACGATCTTGGAGAATCCAG | CAD2222840 | 255 |
| 1860 | GGTCTCATCACCTTTTCTTTATTCGAATACTCCACGA | | |
| 1861 | ggtctcacaagATGCTCAAAAATGAATCCAAACCACC | CAD2218427 | 256 |
| 1862 | ggtctcatcttTCACTCTCTGCTGTTATGATTCTTCCG | | |
| 1863 | ggtctcaccaaATGCTCAAAAATGAATCCAAACCACC | CAD2218427 | 257 |
| 1864 | ggtctcaTCACCTCTCTGCTGTTATGATTCTTCCG | | |
| 1874 | ggtctcacaagATGAACACATTCATCGAGGCAGTCA | ETP9 | 258 |
| 1875 | ggtctcatcttCTAAATCAAGTCTGTGGCAAAGTCA | | |
| 1876 | ggtctcaccaaATGAACACATTCATCGAGGCAGTCA | ETP9 | 259 |
| 1877 | ggtctcatcacAATCAAGTCTGTGGCAAAGTCATCT | | |
| 1945 | ttcatcaaaaaacacagttatccaaATGTTTAGCAGCCCGTTCAA | CAD2216215 | 273 |
| 1946 | tgaacagctcctcgcccttgctcacTTTGAATGTCACCACATGATCTGC | | |
| 1943 | tctcggcatggacgagctgtacaagATGTTTAGCAGCCCGTTCAA | CAD2216215 | 274 |
| 1944 | cactcacgtctctctccccctcttTTATTTGAATGTCACCACATGATCTG | | |
| 776 | GGTCTCGCCAAATGGTGAGCAAGGGCGAGGA | mCherry-SKL | 131 |
| 1142 | GGTCTCGTTACAGTTTGGACTCGTCCATGCCGCCGGTGG | | |
| 1083 | GGTCTCAGTAATAAAAAACATACAAAACAAAAC | 3'-γ-ama FR/ pUMA1467 backbone/5'-γ-ama FR/*hyg*/GAPDH IR | |
| 348 | GGTCTCCTTGGATAACTGTGTTTTTTGATGAA | | |
| 1430 | GGTCTCGCAAGCCTGGACTTCTGTTCGTTG | OCD | 166 |
| 1431 | GGTCTCCTCTTTTACAACTTGGAACCAGACG | | |
| 863 | GGTCTCGTGTGACTGTCAGAAAGCGGC | 5‘-ETP1 FR | 148 |
| 512 | GGTCTCGATGCGTACTTGAATAGCTTTTAATAAT | | |
| 592 | GGTCTCCGCatgaaaaagcctgaactcac | *hyg*/3‘-ETP1 FR | |

| | | | |
|---|---|---|---|
| 595 | GGTCTCCGCACAAATAATGGTTTTGAGGCT | | |
| 590 | GGTCTCGGTGCAGGTCTAGATATCGGATC | pUMA1467 | |
| 862 | GGTCTCGCACAGGTGAGCTCGAATTCA | | |
| 863 | GGTCTCGTGTGACTGTCAGAAAGCGGC | 5‘-ETP1 FR/*neo* | 156 |
| 844 | GGTCTCCTTAGAAGAACTCGTCAAGAAGG | | |
| 524 | GGTCTCCCTAAGGAAAAGATAAATAGACGTTAAAAAAG | 3‘-ETP1 FR | |
| 595 | GGTCTCCGCACAAATAATGGTTTTGAGGCT | | |
| 733 | GGTCTCGTGCGCCAAAGCTGCACTGCA | 5'-ETP2 FR | 092 |
| 734 | GGTCTCGCATTTTGGTGTGTATGATTGTATT | | |
| 845 | GGTCTCGCTAATAATATATATTTATCTCGTTCGGTGTTG | 3‘-ETP2 FR | |
| 846 | GGTCTCGCGGtACGGATCTCATTGCC | | |
| 731 | GGTCTCGACCGCAGGTCTAGATATCGGATC | pUMA1467 | |
| 732 | GGTCTCGCGCAGGTGAGCTCGAATTCA | | |
| 735 | GGTCTCGAATGATTGAACAAGATGGATTGC | *neo* | |
| 844 | GGTCTCCTTAGAAGAACTCGTCAAGAAGG | | |
| 733 | GGTCTCGTGCGCCAAAGCTGCACTGCA | 5'-ETP2 FR | 093 |
| 739 | GGTCTCCTTGGTGTGTATGATTGTATTT | | |
| 740 | GGTCTCCCCAAAatgaaaaagcctgaactcac | *hyg* | |
| 847 | GGTCTCGttgccctcggacgagtgc | | |
| 848 | GGTCTCGgcaaagaaataaTAATATATATTTATCTCGTTCGGTGTT | 3‘-ETP2 FR | |
| 849 | GGTCTCGCGGtACGGATCTCATTGCC | | |
| 871 | GGTCTCGCTGATAATATATATTTATCTCGTTCGGTGTT | 3‘-ETP2 FR/pUMA1467/5‘-ETP2 FR | 094 |
| 872 | GGTCTCGCCATTTTGGTGTGTATGATTGTATTTTTC | | |
| 873 | GGTCTCGATGGCCAAGTTGACCAGT | *phleo* | |
| 874 | GGTCTCGTCAGTCCTGCTCCTCGGC | | |
| 948 | GGTCTCCGCAGGTCTAGATATCGGATC | pUMA1467 | 102 |
| 949 | GGTCTCCCAGGTGAGCTCGAATTCA | | |
| 950 | GGTCTCGCCTGTTGACGAGGACGAGACGG | 5‘-ETP7 FR | |
| 951 | GGTCTCCTCACGGTGGTTTGTTTGT | | |
| 952 | GGTCTCGGTGAATGAAAAAGCCTGAACTCAC | *hyg* | |

| | | | |
|---|---|---|---|
| 953 | GGTCTCGACTTATTTCTTTGCCCTCGGA | | |
| 954 | GGTCTCCAAGTTGAATCACTTTATATACGACAAGG | 3‘-ETP7 FR | |
| 955 | GGTCTCCCTGCCATCACATCCTCGAGGGA | | |
| 950 | GGTCTCGCCTGTTGACGAGGACGAGACGG | 5‘-ETP7 FR | |
| 956 | GGTCTCGTCACGGTGGTTTGTTTGT | | |
| 957 | GGTCTCCGTGAATGGCCAAGTTGACCAGT | *phleo* | 103 |
| 958 | GGTCTCCCAACTCAGTCCTGCTCCTCGGC | | |
| 959 | GGTCTCCGTTGAATCACTTTATATACGACAAGG | 3‘-ETP7 FR | |
| 955 | GGTCTCCCTGCCATCACATCCTCGAGGGA | | |

| Number | 5' RACE primer sequence | Target ETP |
|---|---|---|
| 1275 | CTTGGACGACTTTCCAGGAG | ETP8 |
| 1276 | GAGCTCTAACAGTTTATTGA | ETP8 |
